# Spatially and temporally distinct encoding of muscle and kinematic information in rostral and caudal primary motor cortex

**DOI:** 10.1101/613323

**Authors:** James Kolasinski, Diana C. Dima, David M. A. Mehler, Alice Stephenson, Sara Valadan, Slawomir Kusmia, Holly E. Rossiter

## Abstract

Hand movements are controlled by neuronal networks in primary motor cortex (M1). The organising principle in M1 does not follow an anatomical body map, but rather a distributed representational structure in which motor primitives are combined to produce motor outputs. Both electrophysiological recordings in primates and human imaging data suggest that M1 encodes kinematic features of movements, such as joint position and velocity. However, M1 exhibits well-documented sensory responses to cutaneous and proprioceptive stimuli, raising questions regarding the origins of kinematic motor representations: are they relevant in top-down motor control, or are they an epiphenomenon of bottom-up sensory feedback during movement? Moreover, to what extent is information related to muscle activity encoded in motor cortex? Here we provide evidence for spatially and temporally distinct encoding of kinematic and muscle information in human M1 during the production of a wide variety of naturalistic hand movements. Using a powerful combination of high-field fMRI and MEG, a spatial and temporal multivariate representational similarity analysis revealed encoding of kinematic information from data glove recordings in more caudal regions of M1, over 200ms before movement onset. In contrast, patterns of muscle activity from EMG were encoded in more rostral motor regions later in the cycle of movement. Our spatial and temporal analysis provide compelling evidence that top-down control of dexterous movement engages kinematic representations in caudal regions of M1 prior to movement production; an area with direct cortico-motorneuronal connections. Muscle information encoded more rostrally in M1 was engaged later, suggestive of involvement in bottom-up signalling.

## 1. Main text

Mounting evidence supports the encoding of movements in M1 based on kinematics and synergistic muscle activation, rather than the anatomy of the peripheral musculature^1,2^. Measurements from individual M1 neurons in non-human primates reveal the encoding of multiple kinematic features, such as speed, direction, and position in the same cells in a time-varying manner^3^. The same neuronal populations have been shown to encode instantaneous features during motor execution, as well as the target kinematic end point and upcoming movement trajectory^4,5,6,7^.

In the human brain, evidence of neuronal tuning to multiple kinematic features has been reported during the production of intended movements from M1 microelectrode recordings made in tetraplegic patients^8^. The encoding of kinematic features of hand movements in M1 has also been supported by human imaging studies^9,10,11^. Patterns of fMRI activity in sensorimotor cortex have been shown to mirror the relative differences in the final joint configuration across a range of prehensile movements^12^. Similarly, the representational structure of fMRI activity in M1 during finger flexion is consistent with patterns of finger co-use during naturalistic hand movements^13^.

However, the functional relevance of kinematic encoding in M1 to human motor control remains a fundamental unknown. As well as their role in motor output, M1 neurons exhibit rapid and integrative responses to somatosensory signals^14,15^. Kinematic information is inextricably linked to proprioceptive and tactile signals: specific patterns of movement are associated with specific patterns of sensory feedback. Are kinematic motor representations reported in human M1 functionally relevant in the process of top-down motor control, or an epiphenomenon generated by bottom-up sensory feedback during human movement production?

We addressed this question using a spatiotemporal multivariate representational similarity analysis to ask where in the human brain and when during movement production are the kinematics of human hand movements encoded? This approach combined high-field fMRI and MEG data with kinematic data glove recordings made during a broad repertoire of prehensile and non-prehensile hand movements. Probing recordings of human brain activity with high spatial resolution from fMRI and high temporal resolution from MEG offered a powerful means to identify the location and timing of kinematic information encoding. Together this information was used to dissociate the relevance of kinematic information in M1 to top-down or bottom-up processes in motor control, as well as the relevance of alternative muscle-based or ethological action based models.

Ten right-handed participants performed a range of 26 prehensile and non-prehensile hand movements^17,18^ (Table 1, Video S1) in two fMRI sessions (1.5 hours total fMRI data per participant), two MEG sessions (1.5 hours total MEG data per participant), and a behavioural testing session (35 minutes kinematic data recording). In each session participants wore a right-handed 14-channel fibre optic data glove; kinematic data were recorded through-out all sessions. Electromyography (EMG) data were acquired during MEG sessions to validate the movement onset measures calculated from the data glove.

**Table 1:** Outline of the 26 hand movements used in the motor task. Instructional videos presented in Video S1.

| Hand movements |  |
| --- | --- |
| Abduct fingers | Pinch: thumb and little finger |
| Cylinder Grip | Pinch: thumb and index finger |
| Hook Grip | Pinch: thumb and middle finger |
| Spherical Grip | Pinch: thumb and ring finger |
| Index finger flexion (45°) | Ring finger flexion (45°) |
| Index finger flexion (90°) | Ring finger flexion (90°) |
| Index and middle finger flexion (90°) | Ring and little finger flexion (90°) |
| Index finger and thumb roll | Rock fingers |
| Little finger flexion (45°) | Squeeze: thumb and fingers |
| Little finger flexion (90°) | Abduct thumb |
| Middle finger flexion (45°) | Extend thumb |
| Middle finger flexion (90°) | Flex thumb |
| Middle and ring finger flexion (90°) | Twiddle: thumb and index finger |

To probe the spatial and temporal correspondence between patterns of brain activity and hand kinematics, data glove recordings were used to construct a kinematic model quantifying the similarity of the kinematic signals measured during each of the 26 movements (Figure 1: Top row, Figure S2). The kinematic model quantified the distance between the displacement measures for each movement pair across the 14 channels (Pearson correlation), subject to a Fisher Z-transformation and averaged across the 14 recording channels. The resulting kinematic model exhibits strong split-half and inter-session consistency within participant (Figure S1). A grand average of the kinematic model across sessions and participants was subject to non-classical multidimensional scaling for visualisation of the relative dissimilarity of each movement across two dimensions (Video S2). In both the spatial and temporal representational similarity analysis, the kinematic model was investigated alongside two other models. A muscle based model was constructed from high-density EMG recordings (15 channels) made in an independent cohort of 10 participants performing the same range of hand movements (Figure 1: Bottom row). An additional ethological action model classified movements into precision prehensile, power prehensile, and non-prehensile, based on the notion of ethological maps in primate M1^17,16^ (Figure S17).

**Figure 1:**
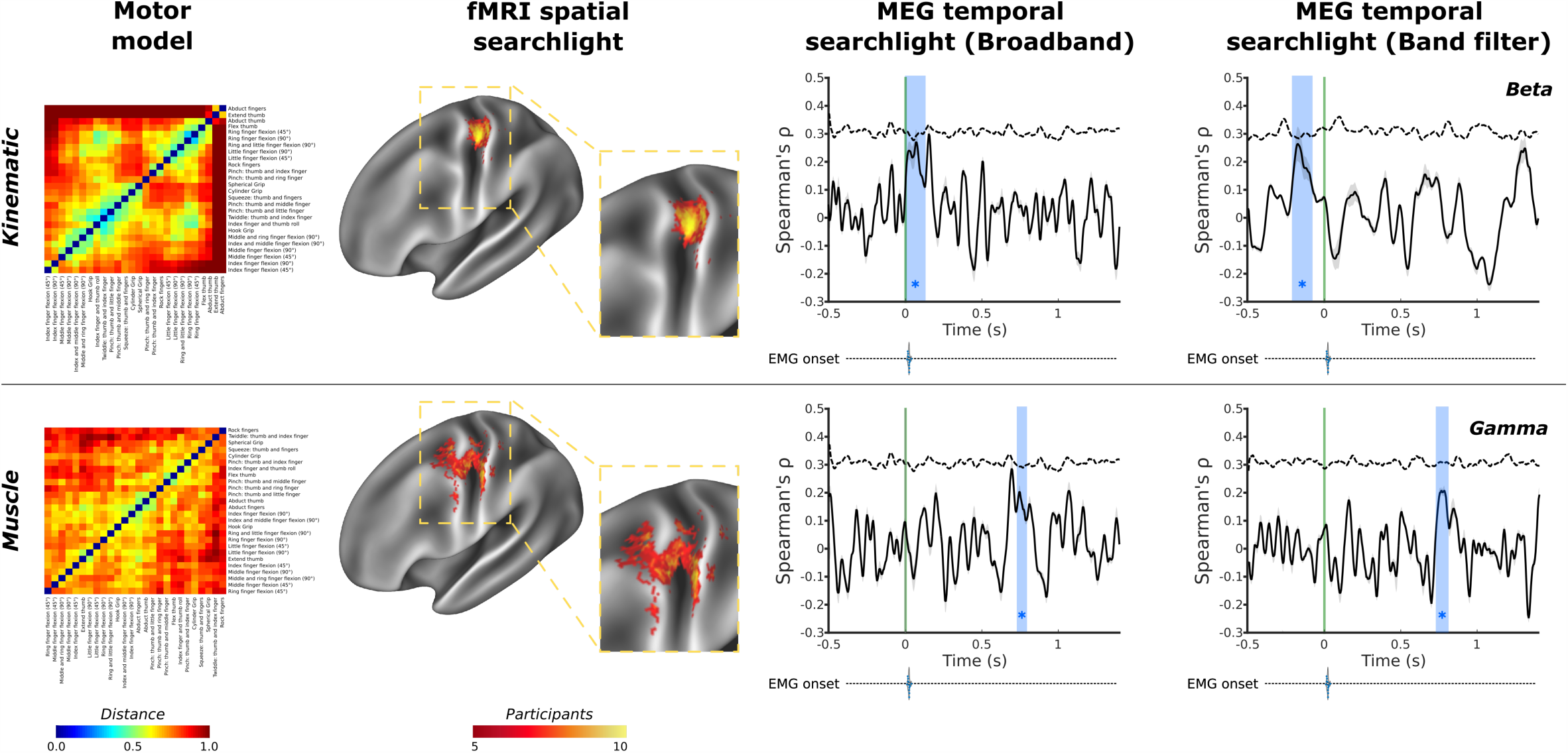
Spatial and temporal evidence for distinct encoding of kinematic and muscle based information in human motor cortex. Kinematic and muscle models of hand movement were used in a spatiotemporal representational similarity analysis. Top row: fMRI data show that kinematic information was encoded consistently in in of primary motor cortex across all ten participants with a peak for all ten participants in Brodmann Areas 4 and 3a; complementary MEG data revealed temporal encoding of kinematic information (blue box) around the point of movement onset in the broadband signal, further decomposition of which revealed encoding prior to movement onset (green line) in the beta frequency, around -210 to -90 ms. The muscle model (bottom row) showed peak consistent encoding in more rostral regions of Brodmann area 4 of primary motor cortex across participants, as well as postcentral regions of Brodmann areas 3b; a temporal searchlight using the muscle model revealed evidence of encoding much later in the cycle of movement around 735-795 ms after movement onset in the broadband signal; further decomposition revealed this encoding of the muscle model originated in the gamma frequency. An ethological action model in line with recent primate studies^16^ was investigated and is presented in Figure S17. Full MEG analysis are presented in Figure 3; green line - movement onset defined by the data glove; blue regions - significant peaks in representational similarity between MEG data and the motor model; dashed line - correlation noise ceiling. EMG onset violin plots based on data presented in Figure S9. Model matrices reproduced in a larger format in Figure S2

We first used high-resolution fMRI data to perform a cross-validated cortical surface-based searchlight representational similarity analysis to find evidence for the spatial encoding of kinematic information during movement. In each participant and each cortical searchlight, the unsmoothed pattern of fMRI activity during movement was used to construct a representational dissimilarity matrix (RDM)^19^. The RDM was compared to kinematic, muscle, or ethological action model, resulting in representational similarity cortical surface maps of Spearman’s *ρ* values for each participant and model. Spearman’s *ρ* surface maps for each model were subject to an omnibus threshold (*α* = 0.01) and used to construct a cross-participant heatmap. This analysis assessed where the relative dissimilarities in the kinematic, muscle, and ethological actions across the different hand movements were mirrored by the relative differences in the pattern of fMRI activity elicited by performing the same movements.

For the kinematic model, the searchlight revealed a strong and very consistent representational similarity in the contralateral pre-central region of the anatomical hand-knob^20^ across participants (Figure 1: top row). Specifically, the fMRI searchlight results revealed the consistent encoding of the kinematic information in Brodmann Area 4 during the production of hand movements across participants (Table 2)^21^. Inspection of the single-subject cortical searchlight results for the kinematic model highlights the consistent and spatially limited correspondence of the kinematic model and fMRI data at the level of individual participants in contralateral M1 (Figure 2A). In the contralateral hemisphere, the peak spatial overlap in the encoding of kinematic information across participants was observed in Brodmann area 4 and 3a; other regions to reach significance at the level of individual participant searchlight analyses, but were not observed consistently across the entire group includes Brodmann Area 3a, Brodmann area 2, 3b, and Brodmann area 6. A highly comparable result was also observed using the kinematic model constructed from the data glove recordings made in the behavioural testing session (Figure S18), highlighting the applicability of this result to real-world hand use in an upright sitting position. No such consistent representational similarity was observed in the corresponding searchlight of movement-related activity in the ipsilateral hemisphere at the group level, however at the level of individual participants significant encoding was observed in greater than three participants included Brodmann areas 4, 3a, and 6 (Figure 2B and Figure S18B).

**Table 2:** Outline of peak anatomical correspondence between movement models and fMRI calculated using across participant cortical heatmaps. Peak regions calculated as centre of gravity of areas of peak overlap; peaks separated by a minimum of 20mm. Vertex positions and anatomical definitions are based on HCP S1200 32k release^21^.

| Model | Peak heatmap overlap (Participants) | Peak Vertex | Anatomical location |
| --- | --- | --- | --- |
| *Kinematic* | 10 | 8053/5378 | Brodmann Area 4 |
| *Muscle* | 10 | 5070<br>8015/8044 | Brodmann area 4<br>Brodmann area 3b |
| *Ethological* | 8 | 8070 | Area 3b |

**Figure 2:**
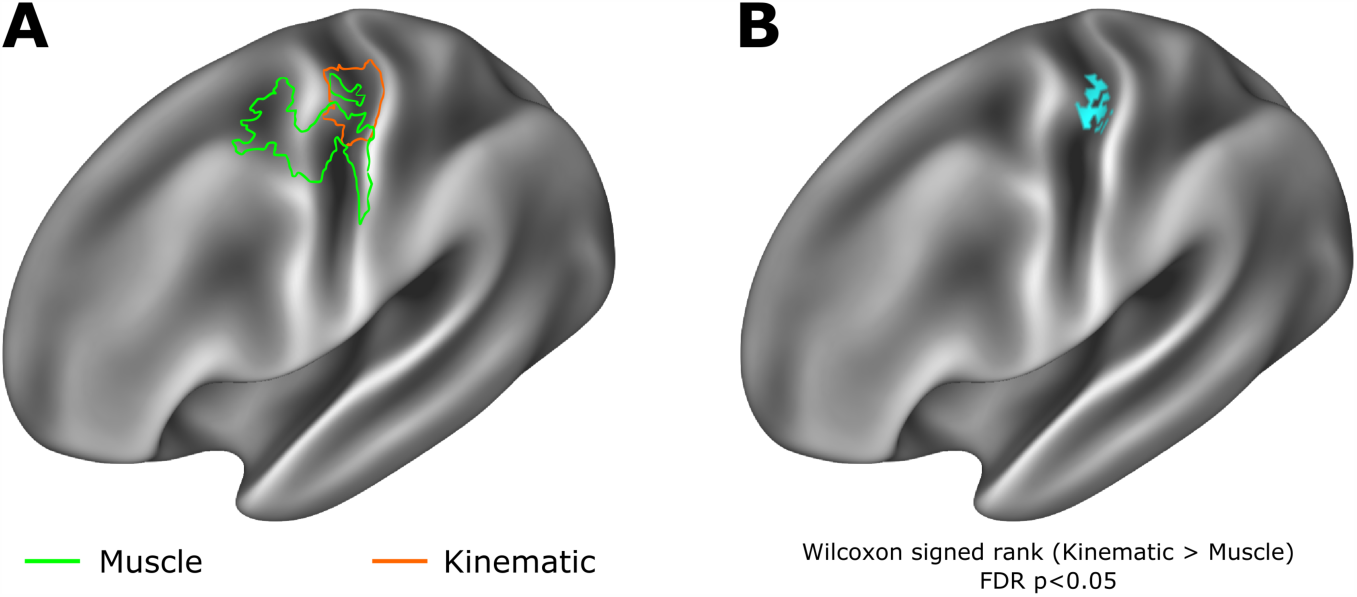
Kinematic and muscle models show evidence of distinct spatial encoding in primary motor cortex. A. Outline of supra-threshold representational similarity analysis results presented in Figure 1 reveal overlapping but distinct encoding of muscle and kinematic information, with muscle information encoding in more rostral regions of Brodmann area 4 and 6, while kinematic information is encoded encode in more caudal regions of primary motor cortex, including Brodmann areas 4 and 3a. (B). A Wilcoxon signed-rank test calculated on Spearmans *ρ* values across the muscle and kinematic spatial searchlights revealed a region at the border of Brodmann area 4 and 3a in which kinematic information showed significantly greater encoding than the muscle model (Statistical maps subject to FDR correction *α* = 0.05).

Equivalent spatial searchlight analyses for the muscle model also revealed supra-threshold activity consistent with encoding in the pre-central region of the anatomical hand knob (Figure 1: bottom row). The muscle model shared representational structure with patterns of brain activity in more rostral and ventral regions compared with the kinematic model, including both areas of Brodmann areas 4 and 6, as well as areas of Brodmann area 3b. This pattern showed less spatial consistency across participants Figures 1 and S6.

In light of the interest in contrasting the kinematic and muscle models^12^, a Wilcoxon signed-rank test (one-sided) was used to compare the vertex-wise *ρ* maps of these two models, which demonstrated the superior fit of the kinematic model in comparison to the muscle model in a localised region principally corresponding to Brodmann Area 4 and 3a^19^ (Figure 3).

**Figure 3:**
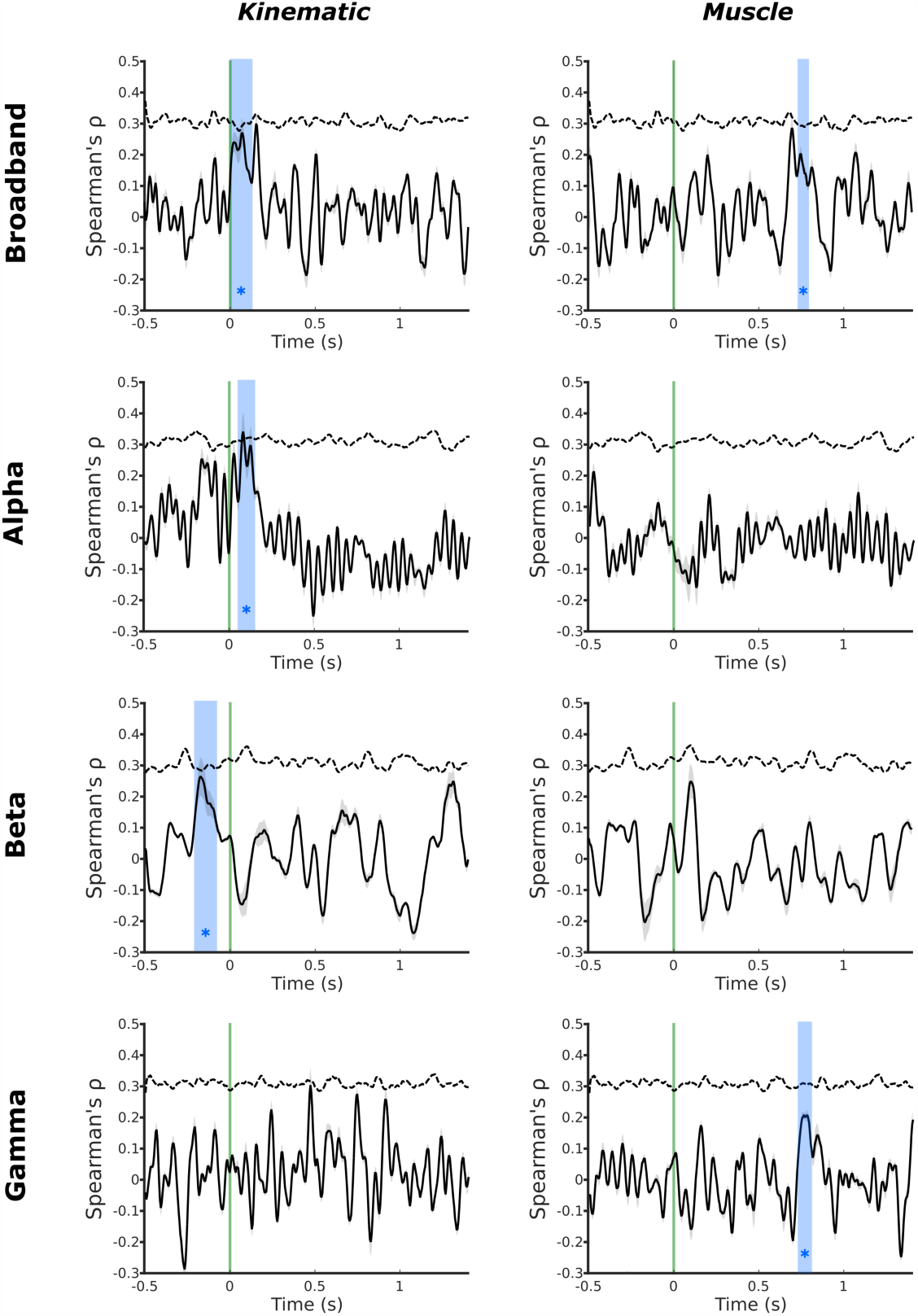
MEG temporal representational similarity analysis searchlight in motor cortex reveals distinct encoding of kinematic and muscle information. Temporal MEG searchlight analysis of the broadband MEG signal revealed encoding of kinematic information around the time of movement onset (5-120ms), contrasted against much later encoding of muscle information 735-785 ms after movement onset. Decomposition of the MEG signal into alpha, beta, and gamma frequencies revealed distinct encoding of the kinematic and muscle models across bands. The kinematic model showed significant encoding in the alpha band after movement onset (55-135 ms) and the beta band prior to movement onset (−210 to -90 ms). In contrast, the muscle model showed significant encoding in the gamma band substantially after movement onset (735 -795 ms). Green line - movement onset defined by the data glove; blue regions - significant peaks in representational similarity between MEG data and the model (1000 shuffled permutations of candidate model RDMs; cluster-forming threshold: *p<* 0.01; maximal cluster distribution (*α* = 0.001)) ; dashed line - correlation noise ceiling.

The ethological action model (Figure S17A) revealed more limited evidence of consistent cortical encoding across participants, centred on somatosensory cortex in the post-central gyrus; specifically Brodmann Area 3b (Figure S17B).

Ultra high field fMRI data analysed at the level of individual subjects offered detailed spatial resolution, revealing spatially distinct encoding of kinematic and muscle information information in the hand knob region of M1. However, fMRI offers relatively poor temporal resolution to understand the point in time at which the kinematic and muscle models match the pattern of brain activity in M1. The boundary between motor and somatosensory cortex is increasingly blurred by evidence of sensory processing in M1^14^ and motor modulation of sensory afferents^22^. The encoding of muscle and kinematic information observed from patterns of fMRI activity may result from top-down control of motor function, or from bottom-up proprioceptive information passed back to M1 and S1. In order to dissociate the driving force behind the spatial model fit observed in the fMRI data, a temporal representational similarity analysis of MEG data was used to identify the point during movement preparation or execution at which kinematic and muscle information is encoded in the M1.

A cross-validated fixed-effects representational similarity analysis was applied, comparing a group average of kinematic and muscle models to the pattern of alpha (7-14 Hz), beta (15-30 Hz), gamma (30-100 Hz) and broad (5-100Hz) band MEG brain activity in M1 (Figure S8) in 20 ms sliding windows during movement preparation and execution. The ethological action model was assessed in equivalent analysis. In light of the interest in contrasting the kinematic and muscle models, the kinematic and muscle models were assessed using a Spearman’s correlation, as well as in a partial correlation to discount the contribution of the other (Figure S20).

Temporal MEG searchlight analysis using the broadband MEG signal revealed encoding of kinematic information around the time of movement onset (5-120 ms relative to movement onset) (Figure 3: top row). Decomposition of the MEG signal into alpha, beta, and gamma frequencies revealed distinct encoding of the kinematic and muscle models across bands. The kinematic model showed significant encoding in the alpha band after movement onset (55-135 ms). In the beta band, the kinematic model mirrored the pattern of brain activity in a significant peak from before movement onset (−205 to -90 ms). In contrast, the muscle model showed significant encoding in the broadband analysis substantially after movement onset (735 - 785 ms), which originated from a temporal correspondence with information encoded in the gamma band (735 - 795 ms relative to movement onset) (Figure 3).

An analogous MEG temporal searchlight analysis during action observation revealed evidence of a correspondence between the kinematic model and brain activity during the movement videos preceding each movement block (Figure S4). During action observation a correspondence between the MEG signal and kinematic model was observed from 220-255 ms and 890-955 ms in the alpha band, 705-735 ms in the beta band, and 545-560 ms in the gamma band, relative to stimulus onset. No peaks in any frequency band were observed for the muscle model or the ethological action model during the period of action observation.

Taken together, the MEG and fMRI results presented here strongly implicate the distinct spatial and temporal encoding of kinematic and muscle information in M1. Specifically, fMRI data suggest kinematic information is represented more caudally in M1, in Brodmann area 4 and 3a, while complementary MEG data suggested kinematic information is encoded prior to and immediately following movement onset in alpha and beta frequencies (Figure 3). In other words, the relative differences in the kinematic structure of a range of different hand movements is encoded in M1 up to 210 ms before the onset of movement can be detected in the hand. In contrast, the muscle based movement model was encoded in more caudal regions of M1, including Brodmann areas 4 and 6 (Figures 1 and 3). Temporally, the muscle model was encoded much later in the cycle of movement, starting at 735 ms after movement onset in the gamma frequency (Figure 3).

These results present strong new evidence in our understanding of movement encoding in M1: suggesting that kinematic features of movements are encoded immediately prior to and around the start of movement, consistent with a role for this organisation’s structure in top-down motor control, while muscle-based organisation was observed in more anterior motor cortex; this information structure was evident in patterns of brain activity at a time suggestive of a role in bottom up signalling later during movement production.

The observation of distinct rostral and caudal representational structures in human M1 is in keeping with an extensive primate literature reporting markedly distinct connectivity profiles along this axis of M1 in non-human primates. Specifically, retrograde labelling studies have reported that the evolutionarily newer caudal region of M1 contains a very high density of cortico-motorneuronal cells (CM cells): those which make monosynaptic connections with motoneurons and are associated with highly skilled movements. In contrast the evolutionarily older rostral M1 contains few, if any, CM cells, relying instead on integrative processes mediated via connections to interneurons in the spinal intermediate zone^23^.

Our current observation of kinematic information encoding in caudal M1 is in keeping with the notion of this cortical region containing CM cells that facilitate specific muscle synergies^24^. The evolutionary development of this caudal M1 region has been specifically associated with the rise of manual dexterity in non-human primates: for example, the existence of large populations of CM cells with monosynaptic connections to motoneurons in the ventral horn of the spinal cord is a hallmark of the ability for independent finger use in the cebus monkey when compared to the squirrel monkey, which has a similar hand structure, but lacks direct cortico-motoneuronal projections^25^. These direct connections via CM cells are not present at birth, but rather develop during early life, and mirror patterns of enhanced dexterous function during infancy and childhood^26^.

In contrast to encoding of kinematic information in caudal M1, we observed encoding of muscle information in more rostral regions of M1 (Figure 3). Lacking CM cells, rostral M1 has been associated with movement via pattern generators or motor primitives via connections to spinal interneurons. In cats, which exhibit only a rostral M1, electrical stimulation to motor cortex elicits movements restricted to very precise muscular anatomy^27^, rather than the patterns of complex movement observed in similar studies of non-human primates^16^. In addition, the inputs to rostral M1 differ from caudal M1: neurons responsive to deep muscle or joint sensory input are concentrated in rostral M1, while cutaneous sensory inputs are concentrated in caudal M1^28,29,30^. Our results provide functional evidence for organisational and temporal differences in the previously described rostral and caudal divisions of M1. Caudal M1, with its direct motoneuronal projections, here showed evidence of encoding movement kinematics, prior to and immediately following movement onset, during the production hand movements. Rostral M1, with its strong deep muscle/joint sensory inputs, showed evidence for the encoding of muscle-based information derived from EMG recordings, which occurred 735-795ms after movement onset, strongly consistent with bottom up sensory signalling from deep joint and muscle receptors. This spatial and temporal dissociation of functional organisation in M1 provides a unique insight into the cortical control of dexterous movements.

Information contained in the kinematic model showed temporally distinct correspondence to information contained in the alpha and beta bands of the MEG data. From 210 ms to 90 ms before movement is detected, the representational structure in the M1 beta band corresponds significantly to the representational similarity of the kinematics of the upcoming movement. The correspondence between the kinematic model and the information contained in the beta frequency band is consistent with the broad literature concerning the role of this oscillatory frequency in motor control. Beta oscillations are observed at rest; it is well established that beta activity is suppressed immediately prior to and during movement: movement-related beta desynchronisation (MRBD), and then rebounds following movement cessation: post-movement beta rebound (PMBR)^31^. The magnitude of the reduction in beta-band power observed prior to movement onset in motor cortex has been shown previously to relate to the degree of uncertainty in the upcoming movement^32^ or action anticipation^33^. Previous comparisons of beta desychronisation made across kinematic and kinetic tasks concur: the strength of MRBD is correlated with the physical kinematic displacement of a given hand movement rather than the magnitude of muscle contraction^34^. Similar patterns of desynchronisation are observed in alpha band activity, where ERD in M1 corresponds to increased activation in the region^31^, with post-motion event related synchronisation in M1^35^. Here we demonstrate that there is a link between information contained in the beta frequency in M1 before movement onset and the subsequent kinematics of hand movements (Figures 1 and 3), suggesting that important information about the upcoming motor command may be encoded within these oscillations^34,36^.

The post-movement peak in kinematic information encoding in the alpha band was observed early after movement onset, during a window of time in which the magnitude of ERD continues to increase after movement has begun^37^.

The observed concurrence between the muscle model and patterns of brain activity measured by MEG occurred some time after movement onset (735-795 ms, Figure 3). An increase in the amplitude of gamma oscillations has previously been reported during motor execution: movement-related gamma synchronisation (MRGS)^38,39^. Increased gamma frequency power is correlated with the size of a given movement, but their strength does not persist during isometric contraction. However, increases in gamma power in M1 are not observed in passive movement conditions, suggesting that gamma activity is not directly associated with muscle activity alone, but rather muscle activity associated with limb movement in combination with the associated sensory feedback^40^. The observed pattern of muscle information encoding in the gamma frequency after movement onset in this study is therefore in keeping with known temporal patterns of MRGS in M1 during movement.

Hand kinematics have previously been investigated in the context of human fMRI. Relative differences in target joint position at the end of a hand movement have been shown previously to mirror the relative differences in the fMRI signal in a broad region of sensorimotor cortex^12^. Additional work considering unidigit and multidigit flexion has demonstrated that patterns of M1 fMRI activity associated with such movements are better explained by kinematic models of digit co-use than by competing muscle-based models^13^. In the present study we have used MEG and 7T BOLD fMRI to fundamentally extend on these findings. Specifically in the context of fMRI, high spatial resolution fMRI data enabled us to reveal a spatial dissociation in muscle and kinematic information encoding in M1 along the rostro-caudal axis (Figure 3). Specifically, we have been able to pinpoint a region of caudal Brodmann area 4 in which kinematic information shows significantly greater encoding than muscle information^20^. Taken alongside evidence from MEG for a temporal dissociation of kinematic and muscle information during the movement cycle, these data strongly implicate kinematic organisation structure in top-down control of hand movements.

The fMRI spatial searchlight analysis did not reveal evidence of consistent encoding of kinematic information in ipsilateral M1 across participants (Figure 2). Previous fMRI studies provide evidence for the activation of ipsilateral M1 during the production of individual uni-digit movements^41,42^ but not multi-digit sequences of uni-digit movements^43^. The present study considered a broad array of naturalistic hand movements, engaging a wide variety of hand kinematics, involving simultaneous and/or sequential movement of different digits. It is possible that unlike sequences of uni-digit movement, these more complex movements do not drive the circuits of ipsilateral M1 as uni-digit movements do^41,42^.

Previous studies have made direct comparisons between muscle-based models and kinematic models, arguing for the latter as an organising principle in the encoding of hand movements^13,12^. As with previous studies, the present findings do not rule out the existence of muscle representations in M1, but rather support the existence of highly organised muscle representations structured around movement kinematics rather than muscle anatomy. The assertion perhaps explains the fractures and repetitions observed in muscle representations during the search for an M1 body map^44^.

The ethological action model reported less consistent patterns of fMRI encoding, centred on the postcentral gyrus, consistent with activation in S1 (Figure S17). The ethological action model also did not reveal any significant peak in the temporal representational analysis. It is possible that while at a coarse level, ethological maps exist in the primate cortex, the concept of ethological organisation does not extend down to the fine-grain level of individual encoding of human hand movements; in other words, the broad motor reportoire of the human hand may not be encoded on the basis of the functional role of each movement. However, in the case of the primate, the coarser division of movements based on the functional role of the entire upper limb, including the hand (e.g. feeding, reaching), may play a role in the way the cortex is organised^45^. The observed patterns of post-central activity may alternatively result from selective disinhibition of S1 by M1 during motor activity, though such direct cortico-cortical signalling remains speculative in the human brain^22,46,47^.

Analysis of the action observation period of the MEG data preceding each movement block also provided some support for the kinematic encoding of information in M1 (Figure S4). Previous MEG data acquired during action observation have demonstrated characteristic changes in M1 activity comparable to action execution^48^. Analyses of event related desynchronisation (ERD) in M1 during action observation have suggested a peak change in the mu frequency as the observed movement evolves^49^. These observations are potentially consistent with the pattern of kinematic model fit observed in the alpha and beta band MEG data during action observation, when the trajectory of movement has become clear (Figure S4). Additional work considering the encoding of kinematic information in oscillatory alpha band activity in M1 suggests that the observation of stimuli consistent with biological motion is sufficient to induce ERD in this frequency band^50^, potentially consistent with the notion that during observation of biological motion, M1 may encode kinematic information.

The data presented in this study rely on complementary information acquired from BOLD fMRI and MEG, though the remit of this work does not extend to fusion of the two modalities. BOLD fMRI provides only an indirect measure of neuronal activity based on haemodynamic changes associated with the execution of a given task^51^, which can be resolved with a relatively high degree of spatial specificity with 7T imaging. In contrast, MEG reflects a more direct, temporally-rich, measure of neuronal activity. While the origins of the measured signals differ, compelling recent evidence provides non-coincidental data to support the notion of shared information across MEG and fMRI measures of brain activity across a wide range of frequency bands^52^; similar correspondences have been reported from invasive electrocorticography data^53^. However, the spatial component of MEG data must be inferred from mathematical modelling. Despite advances in the context of MEG source localisation, this feature of MEG analysis limits the spatial specificity of the measured signals, which integrate information across relatively large tissue volumes in comparison with fMRI^54^. It is therefore not possible to definitely co-localise the signals from MEG and fMRI data. Thus the motor cortex MEG signal used in the temporal multivariate searchlight analysis could have been influenced by signals from adjacent somatosensory cortex; mu-rhythm activity has been shown to associate with sensorimotor BOLD activity^55^. However, previous data from comparative MEG/fMRI studies has suggested a broad association of the sensorimotor alpha frequency signal with the BOLD activity in the post-central gyrus, and the beta frequency with BOLD activity in the precentral gyrus^56,57,58,59^, a similar gradient has been supported broadly by intracortical recordings from non-human primates^60,61^. Here we observe a pre-movement encoding of kinematic information in the beta frequency, and a similar peak immediately after movement onset in the alpha frequency (Figure 3). It is therefore possible to speculate that the beta frequency encoding is more likely to represent pre-central activity in motor cortex, which would again support the conclusion that kinematic information is involved in the top-down control of dexterous movement.

In light of the inability to definitively co-localise fMRI and MEG signals, we have harnessed the respective spatial and temporal strengths of these two methods in independent analyses, rather than using the spatial information from the fMRI to directly inform the temporal analysis of the MEG, making assumptions regarding the shared spatial precision of these two methods.

In this work we apply a rich multi-modal design with multivariate analysis to provide evidence for spatial and temporal dissociations of kinematic and muscle-based information in human M1 during hand movement. Mounting evidence for the encoding of complex kinematic information in M1 from this and other work continues to blur the boundary between primary somatosensory and primary motor cortex: even M1 neurons have been shown to rapidly consolidate sensory torque information across multiple joints^15^. The notion of kinematic representation in M1 is compatible with recent evidence of the tight integration of information across the central sulcus^62^, whereby S1 encodes the current body state, while M1 encodes the kinematics necessary to achieve the intended body state. Such a system of motor control would see kinematic information encoded prior to movement onset as a prediction for the future sensory inputs expected by S1 when a movement has been achieved^63^.

## Acknowledgments

J.K. holds a Wellcome Trust Sir Henry Wellcome Postdoctoral Fellowship (204696/Z/16/Z). CUBRIC is supported by a Strategic Award from the Wellcome Trust (104943/Z/14/Z). This study was supported by the UK MEG Partnership Grant (MRC/EPSRC, MR/K005464/1). The authors are grateful to Krish Singh for his advice regarding the MEG analysis and for his comments on the manuscript, and to Yi-Jhong Han for his technical assistance with EMG data acquisition.

## Notes

#### Summary of Updates

Revision of manuscript on the basis of peer reviewer comments.

