## Supplemental methods and figures for "Spatially and temporally distinct encoding of muscle and kinematic information in rostral and caudal primary motor cortex"

### 2. Supplementary Text: Materials and Methods

#### *2.1. Participants and Experimental Design*

All data were acquired according to the local university research ethics committee approval in line with the Declaration of Helsinki (Cardiff University School of Psychology Research Ethics Committee: EC.17.03.14.4874 and EC.17.04.11.4885) All participants provided written informed consent and met local MRI and MEG safety criteria.

Ten right-handed participants were recruited in the main study (Age range: 22-30; Mean age: 24.0; Age SD: 2.8; 5 Female). Participants were not currently taking any psychoactive medications, and were right-handed according to the Edinburgh Handedness Inventory[64]. No participants had a history of any disorder affecting tactile sensory or motor function or any history of neurological illness. Each participant undertook five experimental sessions: two MRI scan sessions, two MEG recording sessions, and one behavioural testing session. All participants undertook the behavioural testing session first; the subsequent order of the fMRI and MEG sessions was counterbalanced, leaving a minimum of two weeks between any one MRI and MEG session to minimise the effects of magnetic noise on the MEG signal[65]. The datasets generated and analysed during the current study are available from the corresponding author on reasonable request.

#### *2.2. Motor task and kinematic data acquisition*

During all sessions participants were engaged in a motor task involving the production of a range of 26 hand movements (Table 1) with the right hand while wearing a fibre-optic kinematic data glove (Data Glove 14 Ultra; Fifth Dimension Technologies: 5DT, Orlando, FL, USA). Kinematic data were acquired across 14 independent fibre optic channels (one proximal and one distal sensor per digit, plus one sensor between each digit pair) at 60 Hz. Flexion, extension, pitch, and roll cause deformation in the fibre optic channels, impacting the transmission of fibre optic signals and generating a quantifiable signal change. The behavioural task using the data glove was implemented in PsychoPy (Version 1.84.20)[66,67] using the Python Computer Graphics Kit (CGkit: cgkit.sourceforge.net) SDK wrapper for the 5DT data glove.

Each recording session was divided into task runs; each task run was composed of blocks of a specific movement; each block comprised individual movement trials; details of the number runs, blocks, and trials are specified for MEG and fMRI sessions respectively below. Instructions were presented on a screen in the testing environment. Each task run contained one block of each of the 26 movement types, ordered using a random-without-replacement selection method. Progressive determination effects were minimised by maximising the range of different conditions in each run; presenting all 26 movements once per run[68]. At the beginning of each movement block, participants were shown a 3 second video of the movement to be produced (Video S1). Participants were cued to produce the movement in question in each subsequent movement trial of the block by an expanding and contracting horizontal bar. In each movement trial the bar began at a fully contracted width, coloured red, indicating that the hand should be static and in a resting flat position. The bar subsequently turned green and began to expand symmetrically at its left and right flanks. Once it reached its maximal width, the bar began to contract back to its original width. Once the bar reached its original contracted width, it turned red, signifying the end of the movement trial. Participants were instructed to pace their movements to coincide with the period of expansion and contraction of the green bar, such that their hand assumed a flat position at the beginning and end of each trial, corresponding to the time that the static red bar was presented. The motor task was conducted in a behavioural testing lab, in the MRI scanner, and in the MEG scanner, as detailed below.

None of the grasping tasks in this study engaged participants with real objects; previous work has differentiated motor activity with or without real objects in anterior intraparietal sulcus, but not primary motor cortex: as such an object-free study design seemed appropriate for a study focusing on M1[69].

#### *2.3. Kinematic recording session*

During the behavioural testing session participants performed five runs of the motor task. Participants were seated at a desk with their right forearm supported on a memory foam mount, while wearing the data glove. Participants viewed instructions presented on a 14 inch laptop display. Each movement block comprised a 3 second video of the movement to be produced, a 1 second preparation period and 8 subsequent movement trials; each comprising 1.6 seconds of movement (green expanding/contracting bar), followed by a 0.8 second rest period (red static bar). The transition of the bar from red to green was defined as the go signal. A break period of up to 15 seconds was permitted between each movement block; participants advanced the task with a key-press using their left hand. Excluding break periods each task run was 10 minutes and 3.2 seconds in duration. The four task runs yielded 33 minutes and 16.8 seconds of kinematic data recording per participant.

#### *2.4. Kinematic movement model*

For each participant kinematic data from the behavioural, MRI, and MEG sessions were each processed in parallel. This yielded a separate kinematic model from each session type for each participant. These models were used in subsequent multivariate fMRI and MEG analysis; they captured the kinematic similarities and differences of the 26 distinct movements under study.

Initially the kinematic data from each session and each movement block were epoched into individual movement trials using the time of onset of the green bars and averaged. The resulting 14 channels of data represented the average pattern of displacement of the hand during a movement trial for a given movement, termed the kinematics of the movement: the motion of the hand without reference to the forces that produce this motion. In order to compare this signature of kinematic activity for each possible pairing of the 26 movements the activity pattern of each of the 14 recording channels was correlated channel-wise using Pearson's correlation coefficient, subject to the Fisher $Z$-transformation, and the resulting values were averaged across channels to yield a single measure of the similarity of kinematics across each movement pair. The resulting value was transformed back into a Pearson's r-value and used to construct a 1-r dissimilarity matrix for each movement pair.

The kinematic dissimilarity matrices were averaged across task runs to yield an average fMRI, MEG, and behavioural kinematic model for the group. The split-half consistency and inter-session consistency of these models is outlined in Figure S1. A grand average across all sessions and participants was computed and subject to hierarchical clustering; this resulting clustering was applied to all visualisations of the kinematic model (Figure 1).

#### *2.5. Muscle model*

An independent EMG dataset was acquired in order to construct a model of hand movement dissimilarity on the basis of muscle activity in the hand. An independent cohort of ten participants (Age range: 20-30; mean age: 25.1; age SD: 3.57; 5 female) undertook a more detailed EMG recording than was feasible during the MEG session, while performing the same 26 hand movements. EMG data were acquired using a Biosemi Active 2 system with a 32 channel headbox (Biosemi B.V. Amsterdam). Muscle activity was recorded using touchproof flat active electrodes. Electrodes 1-15 were placed as labelled in Figure S15 closely matched to previously published montages[13,12], namely: first dorsal interosseus (FDI), dorsal interosseus muscles, abductor digiti minimi, abductor pollicis brevis (APB), adductor pollicis, lumbrical muscles, flexor carpi ulnaris, flexor carpi radialis, flexor digitorum superficialis and flexor digitorum profundus, flexor pollicis longus. Electrode 16 was used to rereference the EMG data in subsequent analysis and was placed on the lateral bony protrusion of the elbow. There were also CMS and DRL electrodes, which served as a ground/reference during recording in the Biosemi software; they were placed on the dorsal aspect of the wrist. The EMG data were recorded at 2048Hz.

The EMG recording sessions mirrored the design and setup of the kinematic recording session outlined above and were informed by previous fMRI MVPA work[13,12]. Five runs were recorded in total, each containing 26 trials (one for each of the movements). The EMG data were processed using Fieldtrip[70]. EMG data were rereferenced to electrode 16, rectified and subject to a bandpass filter (20 Hz and 1000 Hz); and epoched relative to earliest measured muscle onset in any EMG channel using an adaptive threshold (activity duration threshold: 200ms) (Hooman Sedghamiz: Matlab File Exchange: Automatic Activity Detection in Noisy Signals using Hilbert Transform.) This resulted in individual trials of 2.0s in duration. These trials were baselined using the fixation cross window at the start of each trial. EMG trial data were then subject to multivariate noise normalisation by weighting channels in trial by the error covariance across the different channels in order to more accurate quantify the true differences between the muscle activity across different movements[71,72]. As in the construction of the kinematic model, the activity pattern of each of the EMG recording channels was correlated channel-wise using Pearson's correlation coefficient, subject to the Fisher $Z$-transformation, and the resulting values were averaged across channels to yield a single measure of the similarity of kinematics across each movement pair. The resulting value was transformed back into a Pearson's r-value and used to construct a 1-r dissimilarity matrix for each movement pair. An average muscle model across all ten participants' data was generated and used to probe the spatial and temporal encoding of muscle based dissimilarities in the brain using fMRI and MEG (Figures 1 and S2).

#### *2.6. Ethological action movement model*

An alternative ethological action based model was constructed based on more recent evidence of ethological maps in primate M1[16], and therefore categorises movements on the basis of their specific action, namely prehensile movements, sub-categorised into precision grip and power grip, and non-prehensile movements[18] (Figure S17). The ethological action model was subject to hierarchical clustering for visualisation.

#### *2.7. MRI data acquisition*

MR data were acquired using a Siemens 7T Magnetom system (Siemens Healthcare, Erlangen, Germany) with a 32-channel head coil. Blood oxygenation level dependent (BOLD) fMRI was acquired with a T2*-weighted multi-slice gradient echo planar imaging (EPI). True axial slices were positioned for optimal coverage of the left and right anatomical hand knob[20]

(TR/TE: 1500/25 ms, resolution: 1.2 mm isotropic, 22 axial slices, flip angle:90; GRAPPA factor: 2; anterior-posterior phase-encoding direction; 391 measurements). Magnetization prepared rapid gradient echo (MPRAGE) structural MRI data were acquired to facilitate BOLD EPI slice placement and for cortical surface reconstruction (TR/TE: 2200/2.82 ms, isotropic resolution: 1.0 mm, GRAPPA factor = 2). An additional gradient echo BOLD EPI acquisition of 4 volumes was acquired using posterior-anterior phase-encoding direction for distortion correction.

#### *2.8. fMRI behavioural task*

During the fMRI acquisitions participants performed a total of ten runs of the motor task (5 runs per MRI session). Participants were led supine with their right forearm supported against their right hip and their elbow supported by a foam pad, while wearing the data glove. Participants viewed instructions via a mirror mounted on the transmit coil and a projector screen mounted at the end of the bore. Each movement block comprised of a 3 second instruction screen ("Prepare to Move"), a 3 second video of the movement to be produced, and a 1 second further instruction screen ("Move"), followed by 5 movement trials, each comprising 1.6 seconds of movement (green expanding/contracting bar), followed by a 0.4 second rest period (red static bar). Each movement block was 17 seconds. In addition to the movement blocks, 8 rest blocks were included in each task run; rest blocks were of equivalent duration to movement blocks and comprised of a 3 second instruction screen ("Rest"), a 3 second video of a static resting hand, and a 1 second further instruction screen ("Rest"), followed by the same period of expanding and contracting bar visual stimuli as the fMRI movement blocks. Rest blocks were positioned randomly in each run, excluding self-adjacency.

#### 2.9. Structural MRI data preprocessing

MPRAGE data were subject to reorientation, bias-field correction and brain extraction using the FMRIB Software Library (FSL) fsl_anat tool[73,74,75] prior to cortical surface reconstruction using FreeSurfer Version 5.3.0[76,77].

#### 2.10. fMRI data analysis

##### 2.10.1. fMRI preprocessing and general linear modelling

fMRI data were subject to standard preprocessing, including motion correction with MCFLIRT[78], brain extraction using BET[74], and high pass temporal filtering (100 second threshold). fMRI data were not subject to spatial smoothing. All fMRI data were subject to manual independent components analysis denoising[79]. Distortion correction was undertaken using FSL Topup to estimate a fieldmap image for use in FSL FUGUE[80]. Undistorted BOLD EPI data were co-registered with structural MPRAGE data using Boundary-Based-Registration from FMRIB's Linear Registration Tool (FLIRT) implemented in epi_reg[81,78,82]. Example fMRI timeseries from a single voxel located in the anatomical hand knob is presented for four participants on a single session in Figure S14.

For each participant and each fMRI run, fMRI data were analysed using a first-level general linear modelling (GLM) approach implemented in FSL FEAT[75] using the FMRIB Improved Linear Model (FILM) to estimate time series autocorrelation and pre-whiten each voxel. Each of the 26 movements was modelled with a separate boxcar regressor with gamma-HRF convolution and its temporal derivative, giving a total of 52 regressors. Parameter estimates were calculated, contrasting each movement type against the rest condition; these voxel-wise maps and an estimate of the residuals from the GLM were resampled into the respective participants' structural space and used in subsequent representational similarity analysis (RSA).

*2.10.2. fMRI multivariate noise normalisation*

In order to account for the spatial structure of the noise inherent to fMRI data, spatial prewhitening of the parameter estimates from each participant and each fMRI task run was conducted. The residuals (R) from the first-level GLM analysis provided an estimate of data not fit by the model regressors across voxels (V) and time (T), from which a V x V covariance matrix ($\widehat{\Sigma}$) can estimate the noise structure across voxels (Equation (1))[71].

$$\widehat{\Sigma} = \frac{1}{T}\mathbf{R^T R} \tag{1}$$

The noise covariance structure was combined with the voxel-wise parameter estimates (P) for a given movement type (k) to generate a spatially prewhitened parameter estimate ($P_k^*$: Equation (2)):

$$P_k^* = P_k \widehat{\Sigma}^{-\frac{1}{2}} \tag{2}$$

*2.10.3. fMRI surface-based searchlight representational similarity analysis*

A surface-based representational similarity analysis searchlight approach was used to identify regions in which the multivariate pattern of BOLD activity mirrored the kinematic and categorical models. This surface-based analysis constrained the voxels under consideration in each searchlight to the grey matter and prevented the issue of sampling of voxels that span a sulcus in a single searchlight, which is inherent to volumetric approaches[83]. A searchlight was constructed at the centre of each vertex within the individual participants' anatomical cortical surface region corresponding to the field of view of their task fMRI data (Figure S7). Each searchlight had a diameter of 10mm. The region of interest of each searchlight was projected from 2D surface to 3D volumetric space using the Connectome Workbench Tool[80], masked by a FMRIB Automatic Segmentation Tool grey matter map[73] and a mask excluding voxels spanning across sulci in the FreeSurfer reconstruction to improve spatial specificity. Spatially pre-whitened parameter estimates were extracted from the resulting volumetric region corresponding to each searchlight.

*2.10.4. fMRI cross-validated distance measures*

Within each searchlight the similarity between each of the spatially pre-whitened voxel-wise parameter estimates corresponding to each of the 26 different movement types was calculated using a cross-validated approach to avoid the possibility of over-fitting the data[84,85]. In each iteration, the parameter estimate maps from one fMRI task run was assigned to fold A and the parameter estimate maps from the remaining nine task fMRI runs were assigned to fold B; squared Euclidean distances were calculated between all possible pairs of the 26 movement parameter estimate maps across these two folds (Equation (3)). Distance measures were calculated across all possible pairs of cross-validation folds and averaged[71]. The use of spatially pre-whitened parameter estimate combined with the cross-validation approach yielded cross-validated Mahalanobis distance representational dissimilarity matrices (RDMs) comparing each of the activation patterns across all possible pairings of the 26 movements. For example, calculation of the distance between movement k and movement l in one iteration:

$$d^2_{Crossvalidated\,Mahalanobis}\left(P^*_k, P^*_l\right) = \left(P^*_k - P^*_l\right)_A \left(P^*_k - P^*_l\right)^T_B \qquad (3)$$

The correspondence between the fMRI-derived RDM in each searchlight and the candidate kinematic and theoretical models was assessed using a Spearman's rank correlation, with the resulting $\rho$ (rho) value was plotted in each searchlight's central vertex on the cortical surface. For statistical inference a fixed effects randomisation test[19] was applied on the individual participant level: correlations using 10,000 condition-label randomisations were undertaken in each searchlight. From each of the permutations, the spatial peak $\rho$ value (rho) was extracted from across the cortical surface, forming a maximum accuracy distribution from which an omnibus threshold ($\alpha = 0.01$) was extracted. The resulting thresholded $\rho$-value surface maps for each participant were resampled onto the Human Connectome Project 32k surface (S1200.L.pial.MSMAll.32k_fs_LR.surf.gii), binarised and used to form a heatmap corresponding to the spatial distribution of the each model fit across participants. In light of the interest in contrasting the kinematic and muscle models, a comparison of the corresponding unthresholded Spearman's $\rho$ cortical surface maps was undertaken using a Wilcoxon signed-rank test (one-sided), subject to FDR correction ($\alpha = 0.05$) (Figure 2).

#### *2.11. fMRI motion considerations*

Variability in the magnitude of fMRI motion across different movement conditions has the potential to influence the observed pattern of results. The potential for noise induced by participant motion was mitigated in a number of ways. First, all data was subject to ICA denoising to remove any characteristic motion artefacts[79]. Second, the multivariate analysis of fMRI data emoloyed herein used spatial prewhitening of the parameter estimates to account for voxel-wise variability in noise to not downweight voxels with high error variance and to account for noise covariance between voxels[71]. Finally DVARS values were calculated for each fMRI timeseries (D: temporal derivative of time courses, VARS: root mean squares variance over voxels). These values quantify for each frame of an fMRI acquisition the magnitude of signal intensity change in comparison in volume N compared with volume N-1, as per the following formula:

$$DVARS(\Delta I)_i = \sqrt{\left\langle \left[ I_i \left( \overrightarrow{x} \right) - I_{i-1} \overrightarrow{x} \right]^2 \right\rangle} \tag{4}$$

Where $I_i$ is image intensity at locus $\overrightarrow{x}$ on frame i; angle brackets denote the spatial average over the whole brain[86]. DVARS are able to quantify corruption of fMRI acqusition due to head motion. DVARS values were extracted for volumes corresponding to each of the 26 hand movements for all participants; the resulting distribution of DVARS values is presented in Figure S12. The profiles of very limited motion across participants during each session of around 10 minutes in duration also demonstrate high quality data acquisition (Figure S13).

#### *2.12. MEG data acquisition*

MEG signals were measured continuously at 1200Hz during the motor task using a whole-head 275-channel axial gradiometer CTF MEG system (CTF, Vancouver, Canada) located inside a magnetically shielded room. An additional 29 reference channels were recorded for noise cancellation purposes and the primary sensors were analysed as synthetic third-order gradiometers[87]. Three electromagnetic coils were placed on three fiduciary locations (nasion, left and right pre-auricular) and their position relative to the MEG sensors were recorded continuously during each experimental block. The head surface and fiducial locations were digitized using an ANT Xensor digitizer (ANT Neuro, Enschede, Netherlands) prior to the MEG recording.

#### *2.13. MEG behavioural task*

During the MEG data acquisitions participants performed a total of ten runs of the motor task (5 runs per MEG session). Participants were sitting upright with their right forearm and elbow supported on a foam armrest, while wearing the data glove. Participants viewed instruction on a back-projected screen in front of them from a projector mounted outside the shielded room. Each movement block comprised of a 2 second period with a central fixation cross, a 3 second video of the movement to be produced, and a 1 second instruction screen ("Prepare to Move") followed by five movement trials, each comprising 1.6 seconds of movement (green expanding/contracting bar), followed by a 0.8 second rest period (red static bar). Each movement block was 18 seconds. The order of movement blocks was randomised within each task run; each movement was presented once per task run.

#### *2.14. Data glove movement onset detection: MEG sessions*

The 14 channels of data glove recordings collected during the MEG sessions were synchronised with the MEG acquisitions. Epoched data glove recordings were subject to onset segmentation using an adaptive threshold (activity duration threshold: 200ms) (Hooman Sedghamiz: Matlab File Exchange: Automatic Activity Detection in Noisy Signals using Hilbert Transform.). A conservative estimate of movement onset was derived by taking the earliest signal onset detected across the fourteen data glove channels for each movement trial. The resulting movement onset time was used to epoch MEG data in further analysis.

#### *2.15. MEG data analysis*

##### *2.15.1. MEG preprocessing*

Each participant's head shape was digitized using Xensor digitizer software (ANT software BV, Enschede, The Netherlands). All MEG analysis was conducted using the Fieldtrip toolbox for EEG/MEG-analysis[70] (Donders Institute for Brain, Cognition and Behaviour, Radboud University

Nijmegen, the Netherlands. See http://www.ru.nl/neuroimaging/fieldtrip). Co-registration was performed in a two stage process: first the fiducial locations were marked on the T1 structural for that participant; the head digitization data was then used to align the data with the MRI, subject to manual adjustment. Alignment was undertaken independently for data from the two MEG sessions.

Data from each movement type were epoched from the 10 task runs and concatenated into a new dataset containing 10 blocks, each containing 5 movement trials. The fixation cross and movement trials were epoched from the overall block. The movement trials were defined relative to the data glove defined movement onset time (movement trial time: 2 s; pre-onset time: 0.5s, post-onset time: 1.5s). The fixation cross period was used as a baseline for the 5 movement trials within each movement block. A high pass filter of 1 Hz and a low pass filter of 100 Hz were applied. MEG analyses were conducted across four frequency bands: alpha (7-14 Hz), beta (15-30 Hz) and gamma (30-100 Hz), and broad band (7-100 Hz). All of the movements trials for a given movement type were concatenated across the 10 task runs, creating a dataset comprising 50 repeats of a movement. At this point the data was visually inspected and those trials containing artefacts were removed from further analysis up to a maximum of 10 trials, such that the minimum number of movements trials per movement included in further analysis was 40.

#### *2.15.2. MEG source reconstruction*

In order to reconstruct oscillatory activity at brain locations directly comparable across participants, the individual anatomical MRI was non-linearly warped to the MNI MRI template. The MNI template was divided into a 10 mm isotropic grid and the inverse of the previously calculated non-linear warp was used to warp the template grid into the anatomical space of each participant. Sensor leadfields were calculated using a semi-realistic volume conduction model based on the individual anatomy[88]. The temporal evolution of source activation at each location in the brain was estimated using a linearly constrained minimum variance (LCMV) beamformer algorithm[89] with the optimal dipole orientation at each voxel estimated using singular value decomposition (SVD). Virtual sensors were then reconstructed from all 3294 voxels by multiplying the sensor level data by the corresponding set of optimised weights. At this stage data were subject to multivariate noise normalization[72,90]: we calculated the error covariance matrix at sensor level and then used this combined with the filters from the LCMV to create the virtual sensor data. This means that sensors with more noise would be down-weighted compared to those with less noise. At this stage the data was also down-sampled to 600 Hz to reduce computational cost.

#### *2.15.3. MEG temporal representational similarity analysis*

The MEG data were split to produce 10 partitions and then averaged within each partition to perform a cross-validated representational similarity analysis to avoid the possibility of over-fitting the data[84,85]. RSA was performed across time using a sliding time window with a width of 20 ms and a time step of 5 ms creating 396 time windows across 2 seconds of the movement trial (0.5s rest; 1.5s movement). After selecting virtual sensors within the left hemisphere motor region of the AAL atlas[91] (Precentral L, 31 sources; Figure S8), the frequency-filtered MEG signal measured during each movement type was compared using a cross-validated approach within each time width. In each iteration, the signals from one MEG data partition were assigned to fold A, and the signals from the remaining nine partitions were assigned to fold B; squared Euclidean distances were calculated between all possible pairs of the 26 signals across the two folds and averaged[71]. The use of multivariate noise normalisation to account for spatial autocorrelation in the MEG signal yielded subject-wise cross-validated Mahalanobis distance RDMs comparing the alpha, beta, or gamma-band signal in the motor ROI across all possible pairings of the 26 movements[72].

Participant-level motor ROI RDMs were averaged in order to perform a fixed-effects analysis. The correspondence between the MEG-derived RDMs and the candidate kinematic and theoretical models across time was assessed using a Spearman's rank correlation, with the resulting $\rho$ (rho) values plotted for each time window. In light of the interest in contrasting the kinematic and muscle models, these were each assessed in a partial correlation to discount the contribution of the other. Randomization testing was used for statistical inference[92], whereby candidate model RDMs were shuffled 1000 times and time-resolved correlation coefficients were recomputed in order to estimate an empirical null distribution. p-values were calculated using a cluster thresholding approach across time. To correct for multiple comparisons, the cluster-forming threshold was set to $P < 0.01$ and clusters in the correlation time-courses corresponding to each candidate model were thresholded against the maximal cluster distribution ($\alpha = 0.001$).

To assess the maximal correlation possible with our data, each participant's RDM was correlated with the average cross-subject RDM; the correlations were then averaged to obtain an upper bound of the noise ceiling[19].

#### *2.15.4. MEG: action observation analysis*

MEG data from the period of action observation during the instruction video preceding each movement block were epoched using the same approach as the MEG data recorded during movement. The fixation cross and action observation trials were epoched from the overall block. The action observation trial was defined relative to the video stimulus onset time (pre-onset time: 0.5s, post-onset time: 3.0s). The fixation cross period was used as a baseline for the action observation period. Temporal representational similarity analyses were conducted using the same apporoach as the MEG movement data, as described above.

#### *2.15.5. MEG motion considerations*

MEG analysis included multivariate noise normalisation to account partially for the effects of motion, where each channels are normalised by an estimate of error covariance across different sensors; this process has been demonstrated to substantially improve multivariate analyses of MEG data[72]. Motion parameters for all MEG acquisitions were extracted and analysed to rule out the possibility of excessive head motion as a potential driving force behind any observed patterns of brain activity. Rotational and translational displacement for each participant and each experimental session are presented in Figure S10. In addition, the motion parameters during each movement block were extracted and the resulting distribution is presented across the 26 different movement types (Figure S11). The profiles of motion across participants demonstrate a high quality data acquisition.

### *2.16. Electromyography with MEG*

Electromyography (EMG) data were acquired simultaneously with MEG data. Three surface EMG electrodes were attached to the right hand underneath the data glove, positioned on abductor pollicis brevis (APB), first dorsal interosseus (FDI) and abductor digiti minimi (ADM). The area under the electrodes was exfoliated and cleaned with alcohol prior to data acquisition. EMG signals were recorded at 1200Hz.

EMG data were initially subject to a bandpass filter (20-1000Hz) and a notch filter (50 Hz). EMG data were epoched and baselined alongside the MEG data. Epoched EMG data were subject to manual artefact rejection. Signals from the three electrodes during each epoch were independently subject to a Hilbert transform and smoothing (5 ms window) prior to activity onset segmentation using an adaptive threshold (activity duration threshold: 200ms) (Hooman Sedghamiz: Matlab File Exchange: Automatic Activity Detection in Noisy Signals using Hilbert Transform.). A conservative estimate of muscle activity onset was derived by taking the earliest signal onset detected across the three EMG channels for each movement trial; any trial in which the onset estimate from the EMG and data glove activity recorded during MEG showed a discrepancy of $> \pm 100ms$ were excluded. Due to constraints of electrode placement alongside the kinematic data glove, measures of activity onset were not robustly measured in all participants. EMG onset data are presented in order to validate the data glove measures of movement onset, which have been used to epoch the MEG data (Figures 1 and S9).

**Video S1: Compilation of instructional videos used at the beginning of movement blocks in all testing sessions.** Movement labels are provided for reference only; labels were not included during the task (VideoS1.mov).

**Video S2: Visualisation of multidimensional scaling of grand average kinematic model constructed across participants and sessions.** (VideoS2.mov).

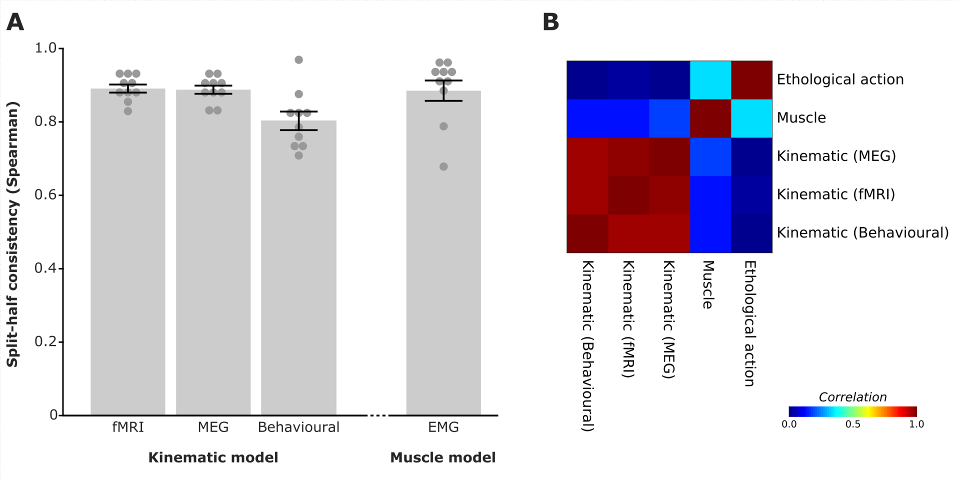


**Figure S1:** (A.) Data-driven kinematic models constructed for each participant and each session type exhibit strong split-half and inter-session consistency. Muscle model reproducibility data also presented. (B.) Strong between-model consistency across kinematic models calculated from data across fMRI, MEG, and behavioural recording sessions; limited shared information across kinematic and muscle models. Ethological action model resulted presented in Figure S17.

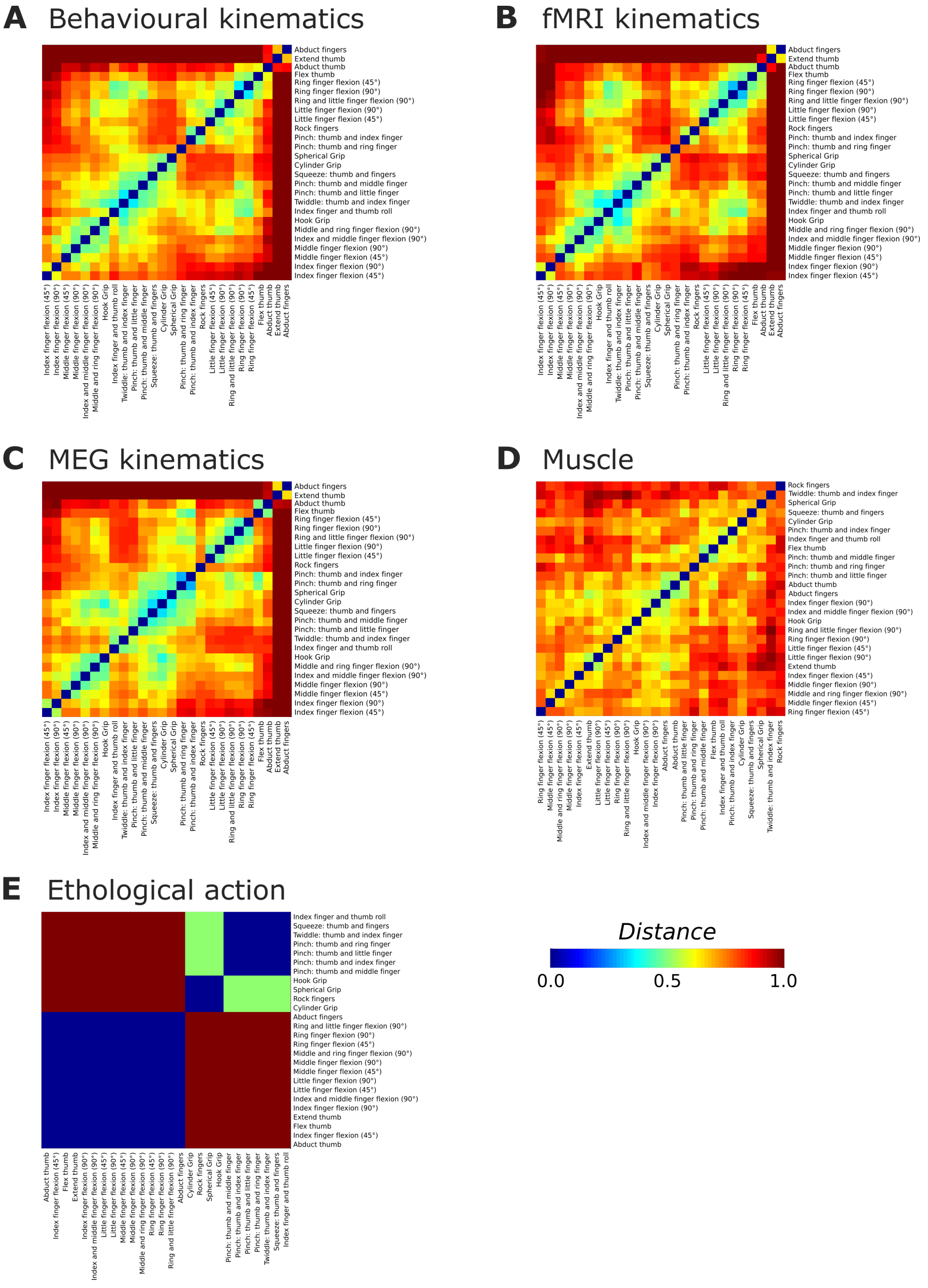


**Figure S2: The average kinematic model across participants for each recording session type, the muscle model derived from an independent cohort of participants, and the categorical ethological action model.**

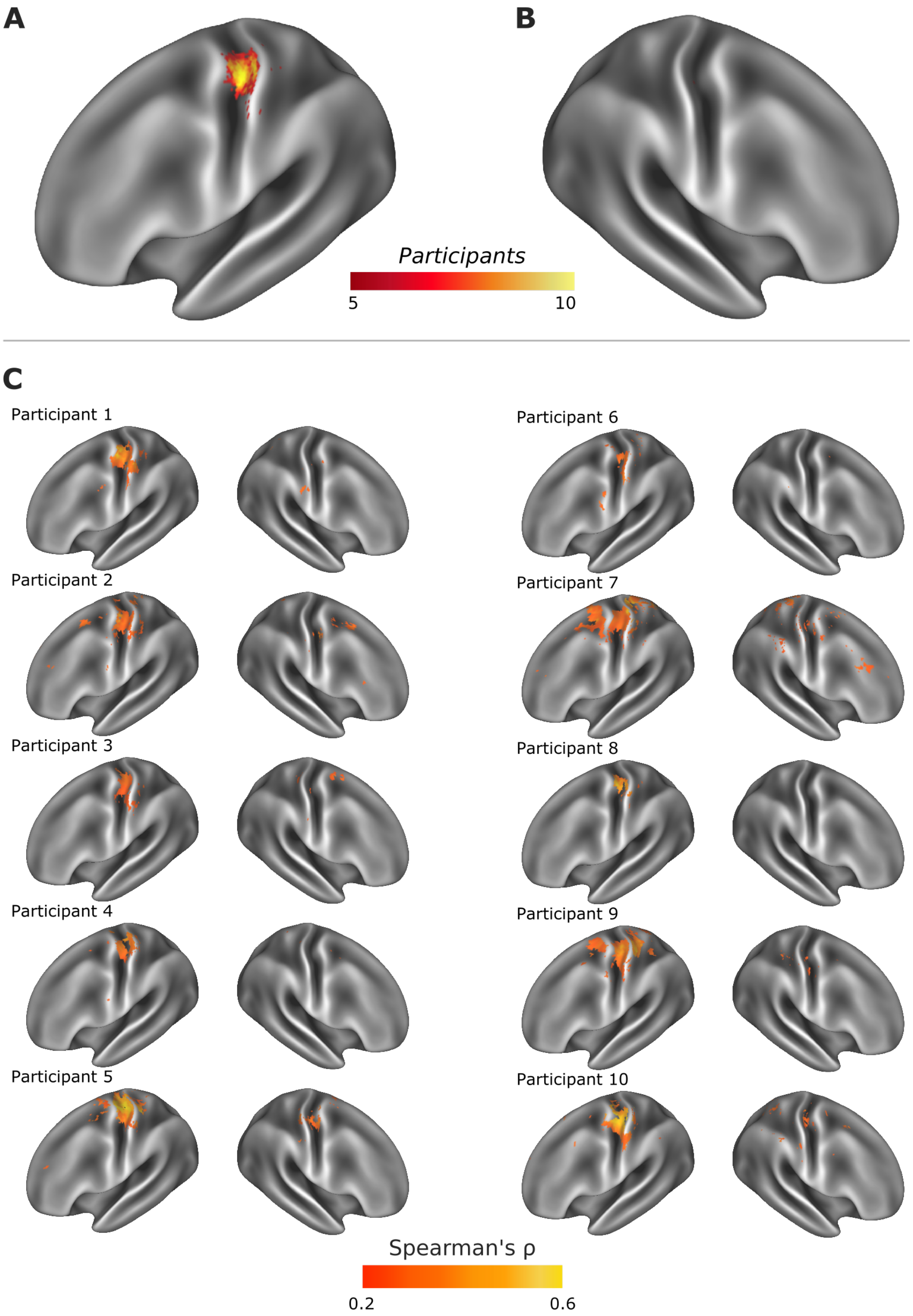


**Figure S3: Single participant fMRI representational similarity analysis cortical searchlights using individual kinematic models of hand movement.** Cortical heatmaps of the left (A) and right (B) hemisphere, showing consistent encoding of kinematic information in the left motor cortex, contralateral to movement. Heatmaps were constructed from individually thresholded cortical searchlights for each participant, derived using their own kinematic model (C) (Omnibus threshold, $\alpha = 0.01$, maximum accuracy distribution calculated from peak correlation value across 10,000 searchlight permutations with label-switching).

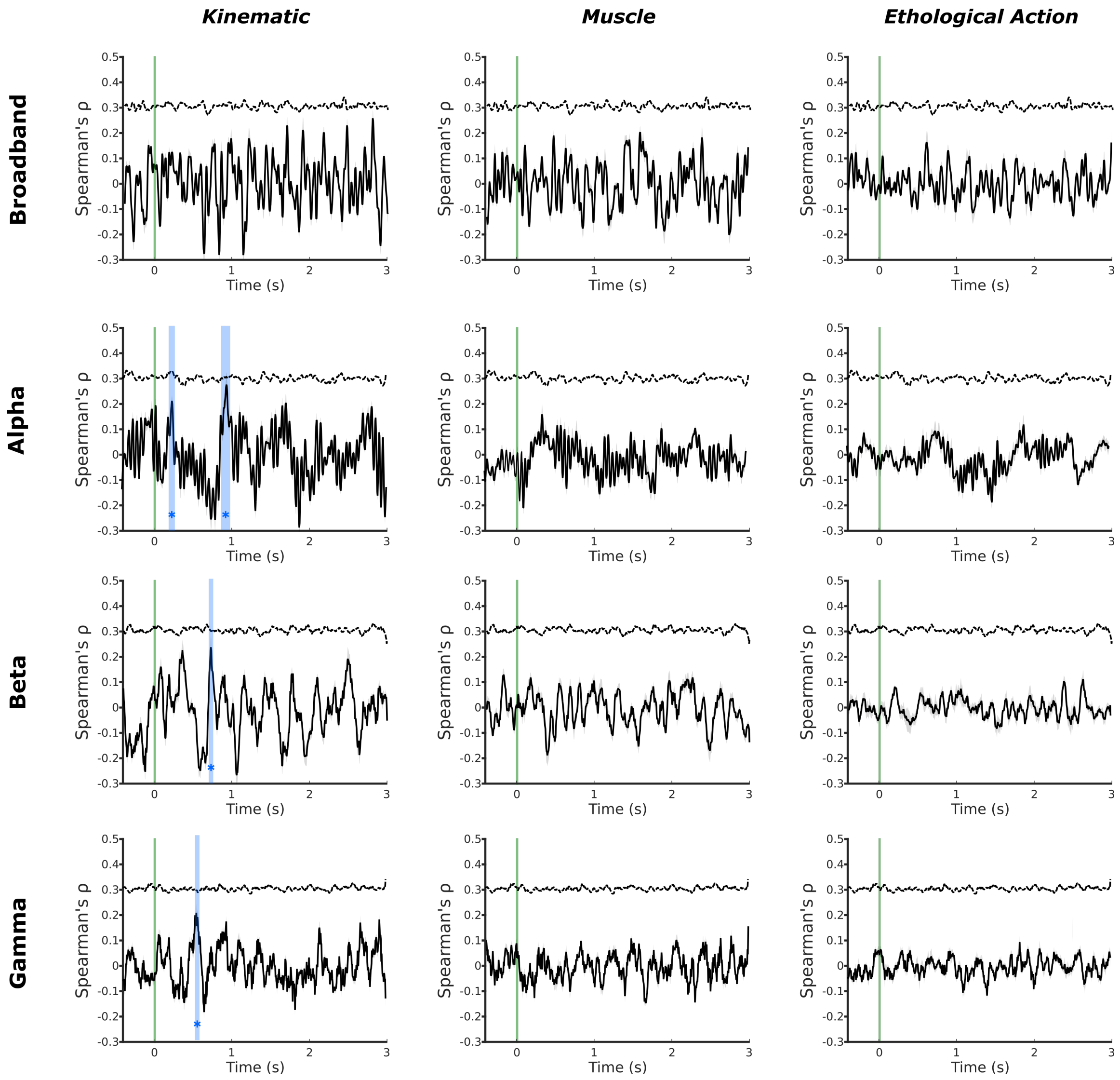


**Figure S4: MEG searchlight analysis during action observation** Evidence of a significant peak in the correspondence between the kinematic model and the alpha frequency MEG signal (220-255ms and 890-955ms), the beta band MEG signal (705-735ms) and the gamma band MEG signal (545-560ms) in the period of action observation. No equivalent concurrence with the MEG signal was observed for the muscle of ethological action model in primary motor cortex during action observation. The green line indicates the onset of the stimulus video; the blue regions indicate significant peaks in representational similarity between MEG data and the motor model; the dashed line indicates noise ceiling. Comparison with MEG temporal searchlight results presented in Figures 1 and 3.

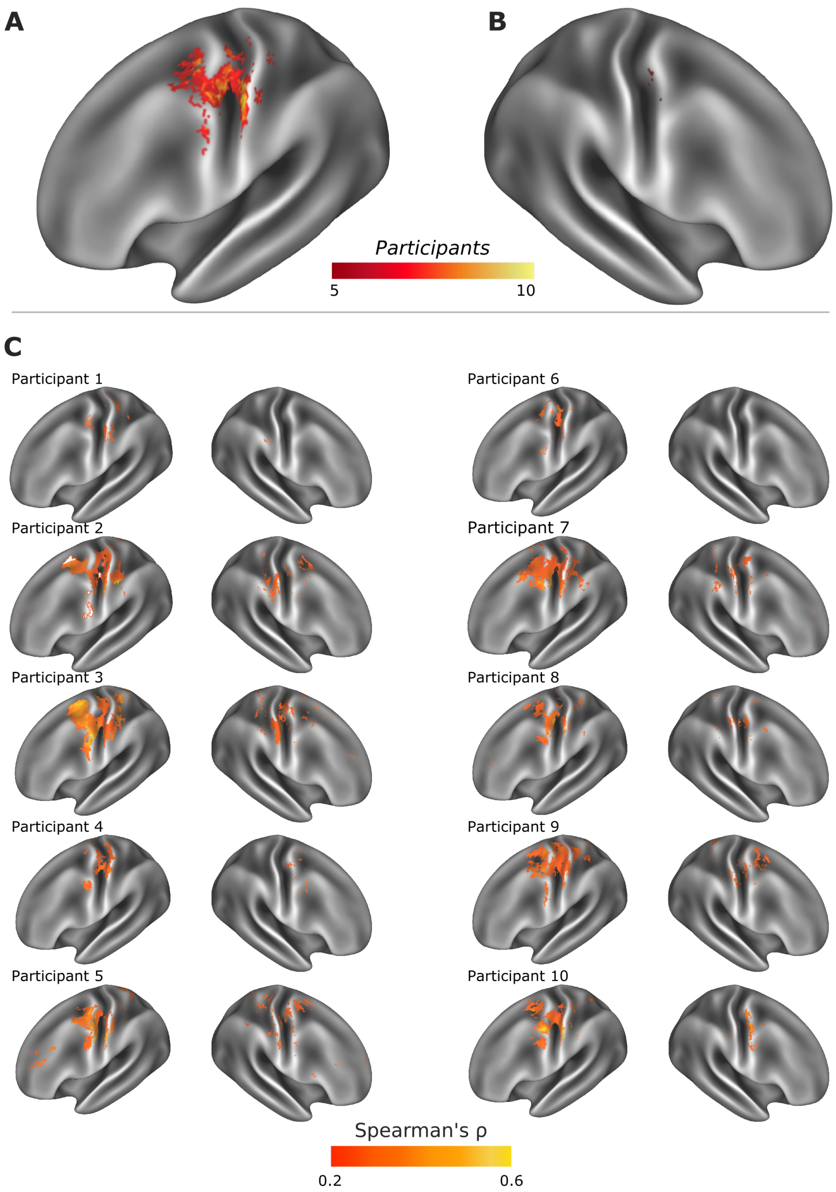


**Figure S5: Individual participant cortical searchlight results using an muscle model of movement encoding.** Cortical heatmaps of the left (A) and right (B) hemisphere, show consistent ncoding of an action model based on EMG recordings in Brodmann areas 4 and 3b. Heatmaps were constructed from individually thresholded cortical searchlights for each participant using a single average muscle model (C) (Omnibus threshold, $\alpha = 0.01$, maximum accuracy distribution calculated from peak correlation value across 10,000 searchlight permutations with label-switching.)

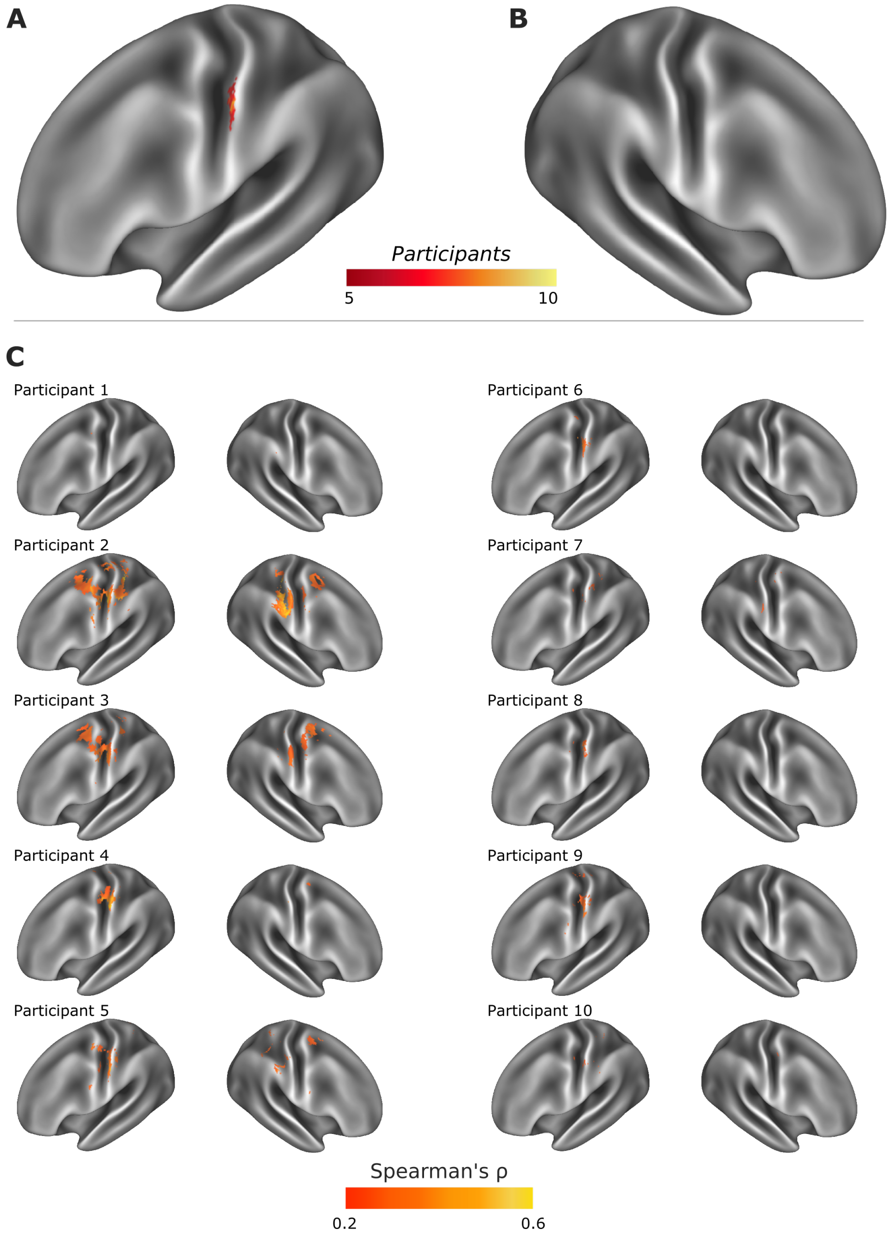


**Figure S6: Individual participant cortical searchlight results using an ethological action model of movement encoding.** Cortical heatmaps of the left (A) and right (B) hemisphere, show limited but consistent encoding of an action model in the left post-central gyrus, contralateral to movement. Heatmaps were constructed from individually thresholded cortical searchlights for each participant using a single categorical action model (C) (Omnibus threshold, $\alpha = 0.01$, maximum accuracy distribution calculated from peak correlation value across 10,000 searchlight permutations with label-switching.)

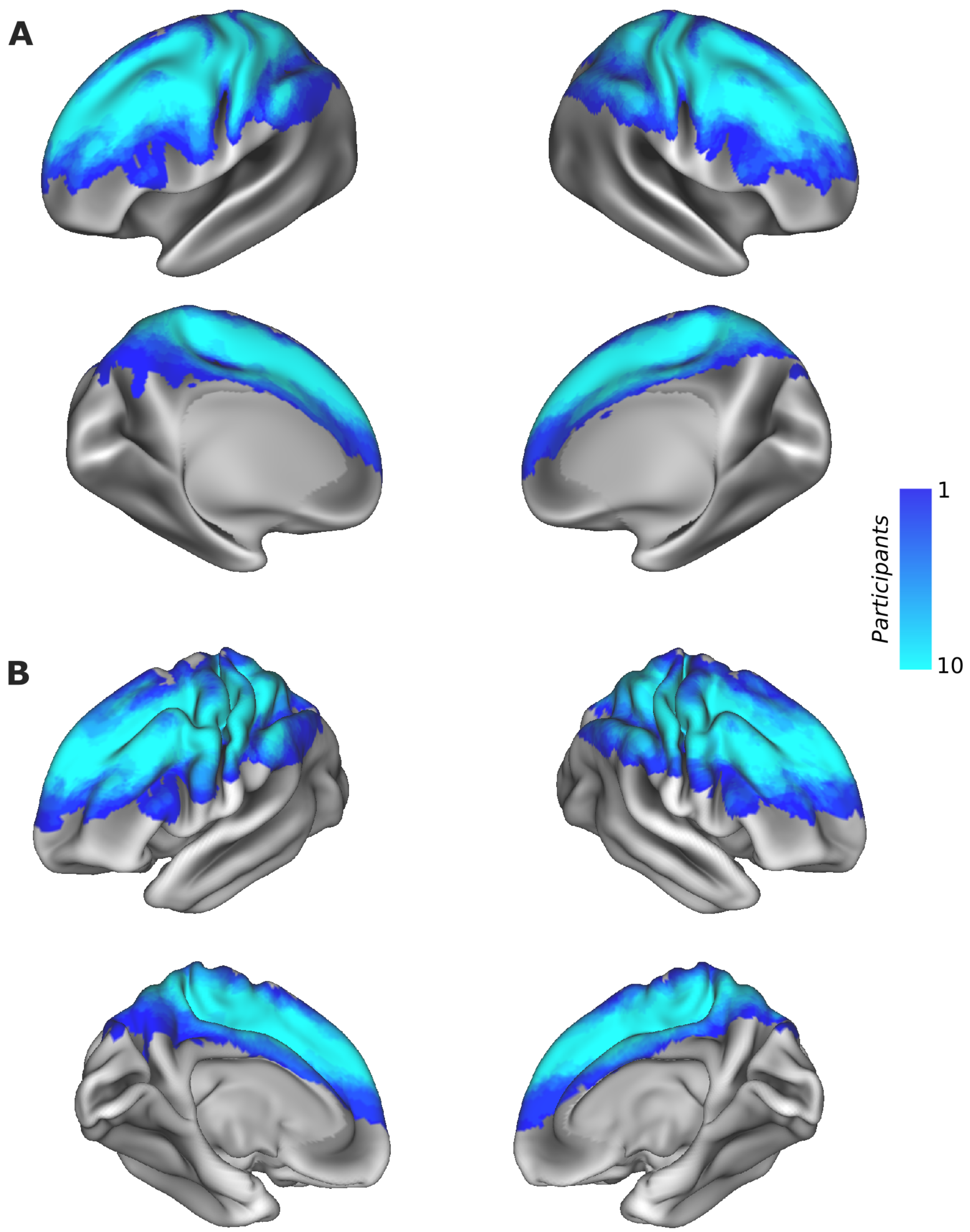


**Figure S7: Heatmap surface visualisation of fMRI data coverage across participants on inflated (A) and midthickness (B) surfaces.**

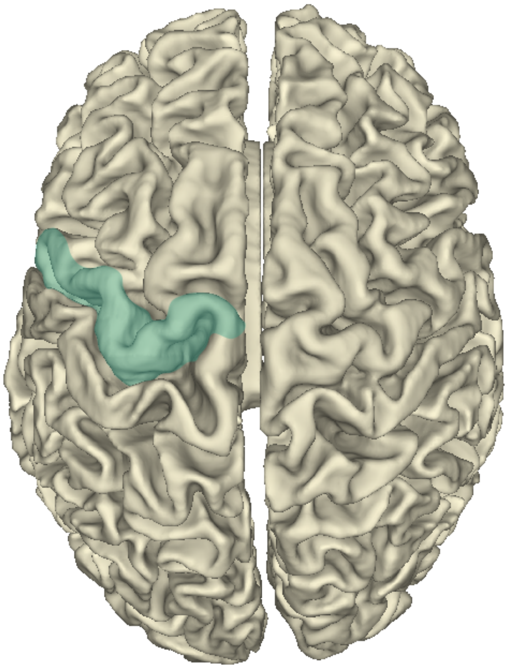

**Figure S8: Surface visualisation of the left hemisphere motor region of the AAL atlas used in MEG temporal searchlight analysis: precentral L.**

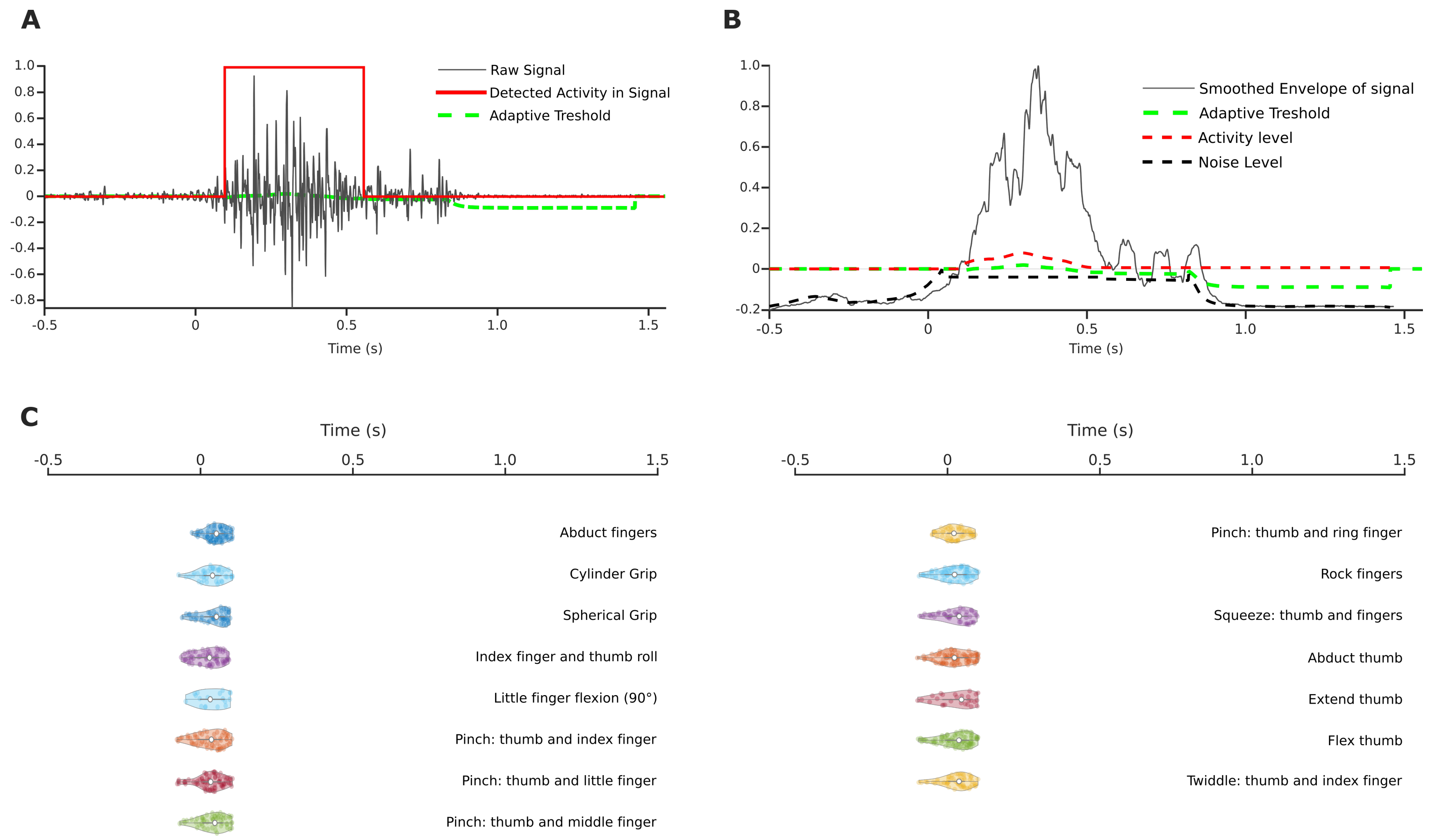


**Figure S9: EMG muscle activity onset time.** A. Example of EMG onset detection using adaptive threshold. B. Smoothed envelope of signal (Hilbert transform) used for automatic onset detection. C. Violin plots visualising the distribution of EMG onset times for trials from which onset could be detected across all participants. Bars represent interquartile range, white dots represent median values which are plotted in Figure 1. EMG data were used to exclude trials where muscle onset time differed from data glove defined movement onset time by $> \pm$ 100 ms.

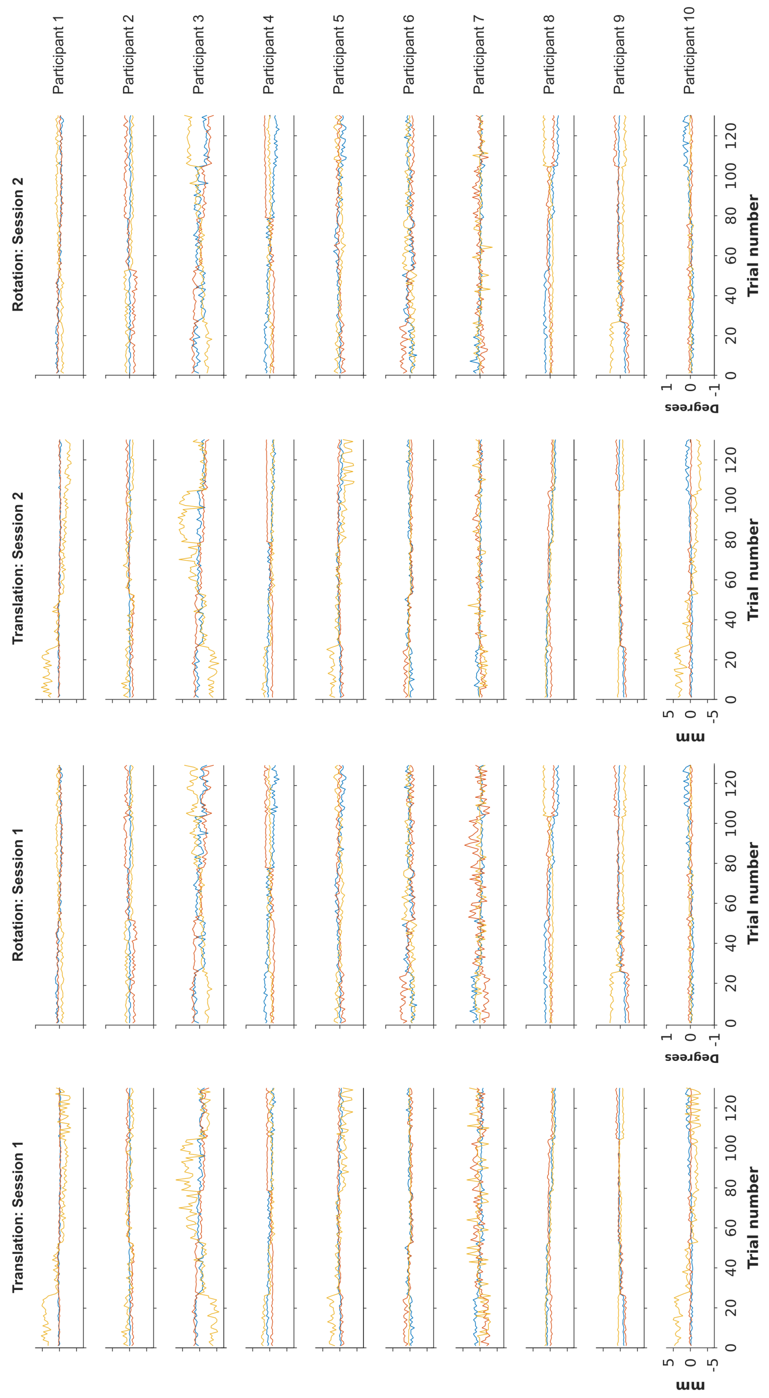


**Figure S10: MEG motion timeseries across participants and sessions. Blue: X-axis, orange: Y-axis, yellow: Z-axis**

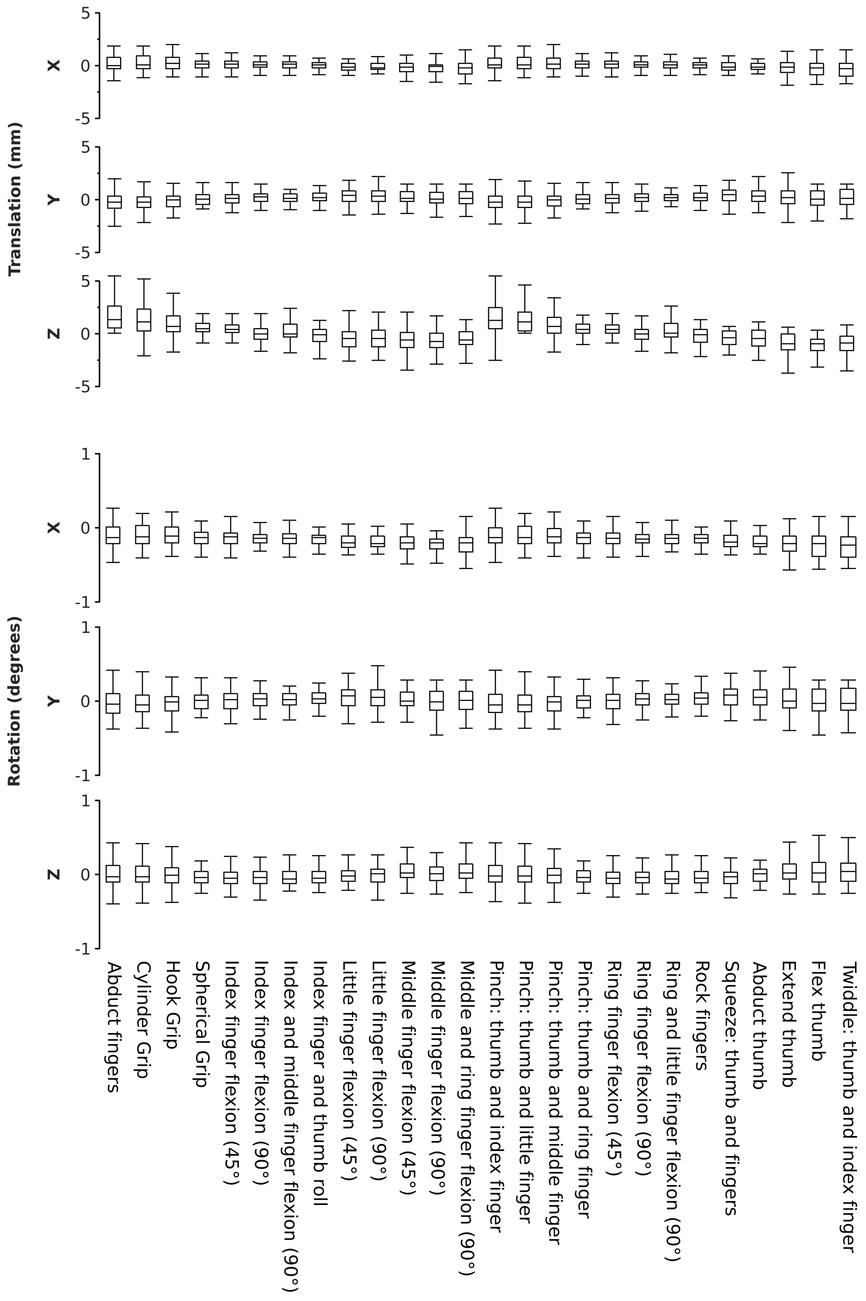


**Figure S11: MEG motion comparison across movement conditions.**

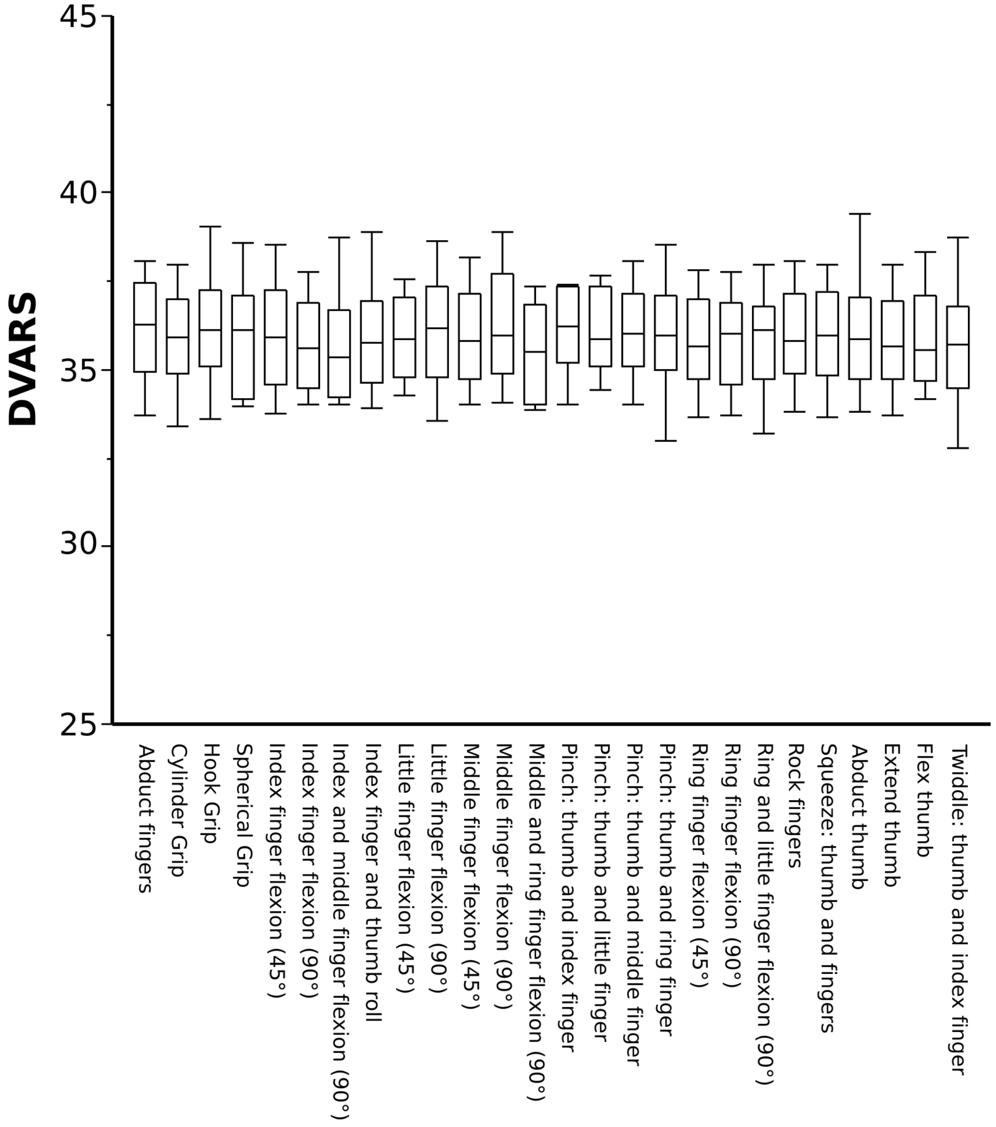


**Figure S12: fMRI motion comparison across movement conditions using DVARS presented in arbitrary units[86,93,94].**

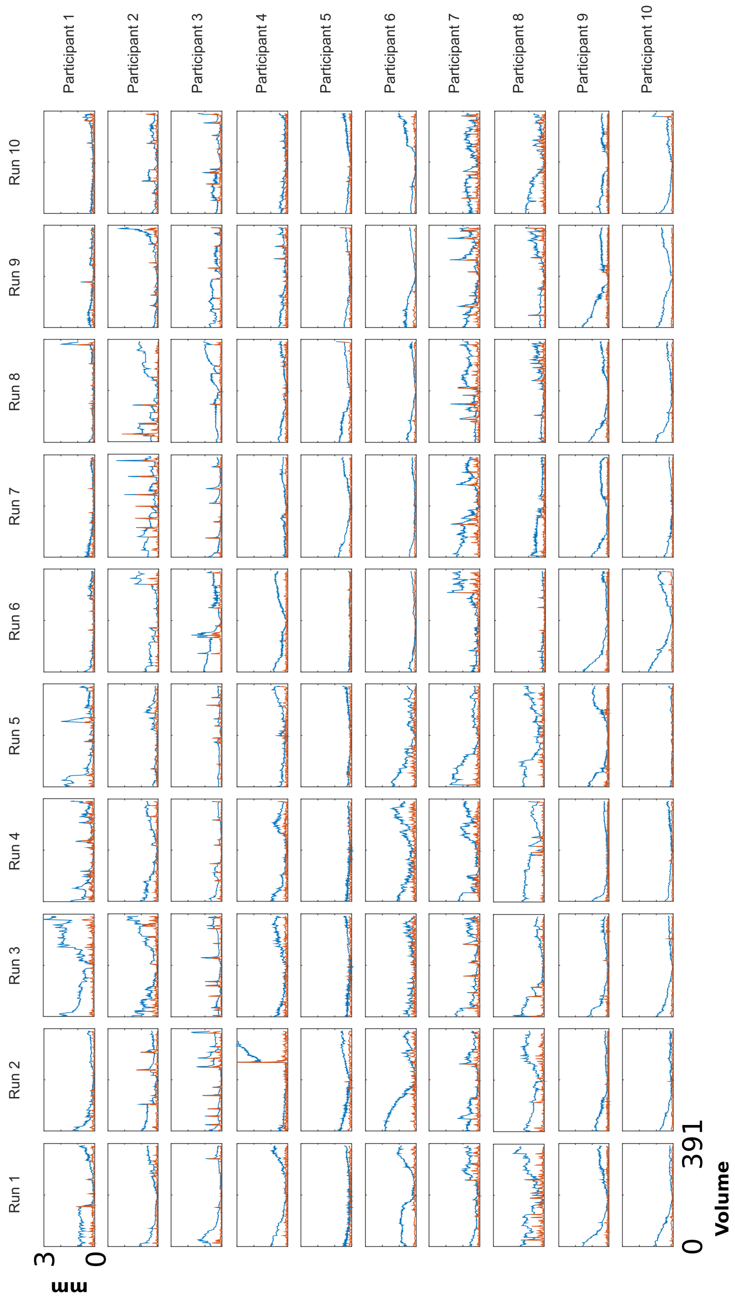


**Figure S13: fMRI motion displacement plots.** Plots of absolute (blue) and relative (orange) motion calculated using FSL MCFLIRT for each participant and each fMRI task run; motion correction was undertaken prior to ICA denoising.

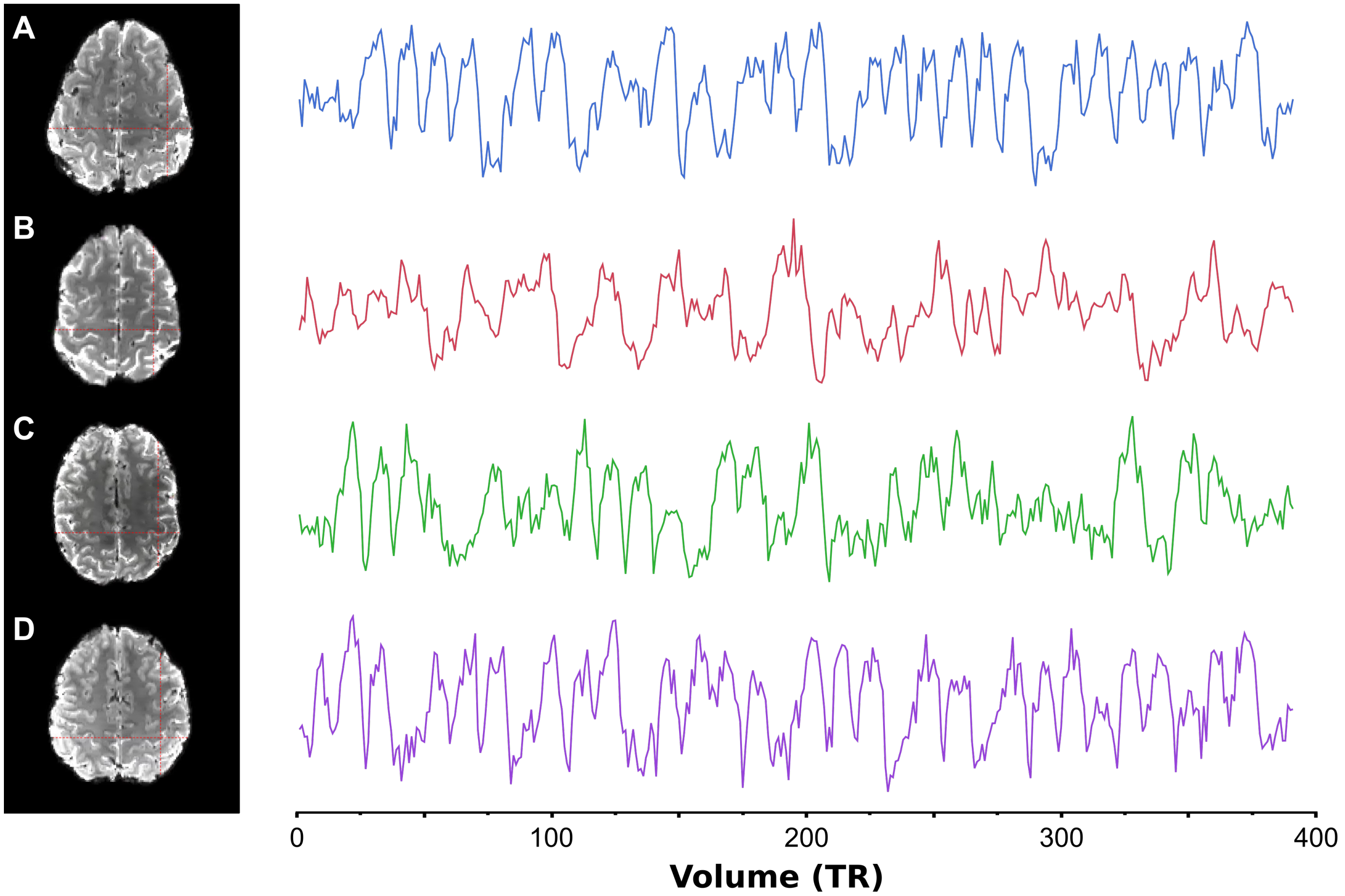


**Figure S14: Example fMRI axial slices and single voxel timeseries from 4 participants.** fMRI timeseries data extracted from a single voxel (red crosshairs) for four participants. Data presented were subject to high-pass filter (100 seconds). fMRI data were not subject to spatial or temporal smoothing.

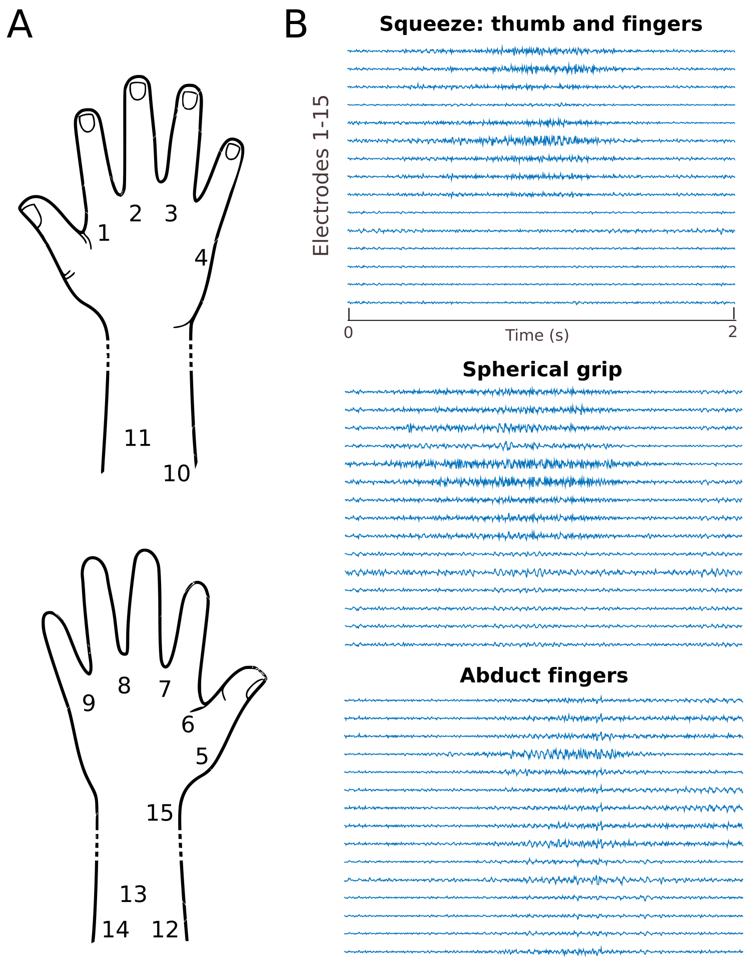


**Figure S15: EMG recordings from an independent cohort used to generate a muscle model of hand movement**. A) Schematic demonstrating electrode placement for EMG recordings on both the palmar and dorsal surface of the hand and forearm covering these muscles: 1: first dorsal interosseus (FDI), 2-3: dorsal interosseus muscles, 4: abductor digiti minimi, 5: abductor pollicis brevis (APB), 6: adductor pollicis, 7-9: lumbrical muscles, 10: flexor carpi ulnaris, 11: flexor carpi radialis, 12-14: flexor digitorum superficialis and flexor digitorum profundus, 15: flexor pollicis longus. Dotted lines indicate region of the forearm compressed for visualisation purposes. This montage replicated that used in previous studies of kinematic and muscle information encoding in M1[13,12]. B) Example EMG trace from one participant for three different movements (squeeze, spherical grip and abduct fingers) showing electrodes 1-15 across the 2s movement time window.

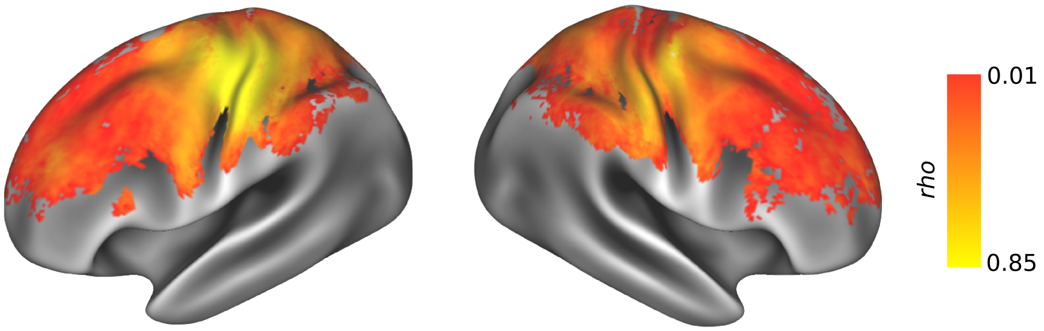


**Figure S16: Noise ceiling calculation for spatial searchlight using fMRI data**. To assess the spatial consistency in RDMs calculated from fMRI data at each vertex, each participant's RDM was correlated with the average cross-subject RDM; the correlations were then averaged to obtain a vertex-wise upper bound of the noise ceiling.

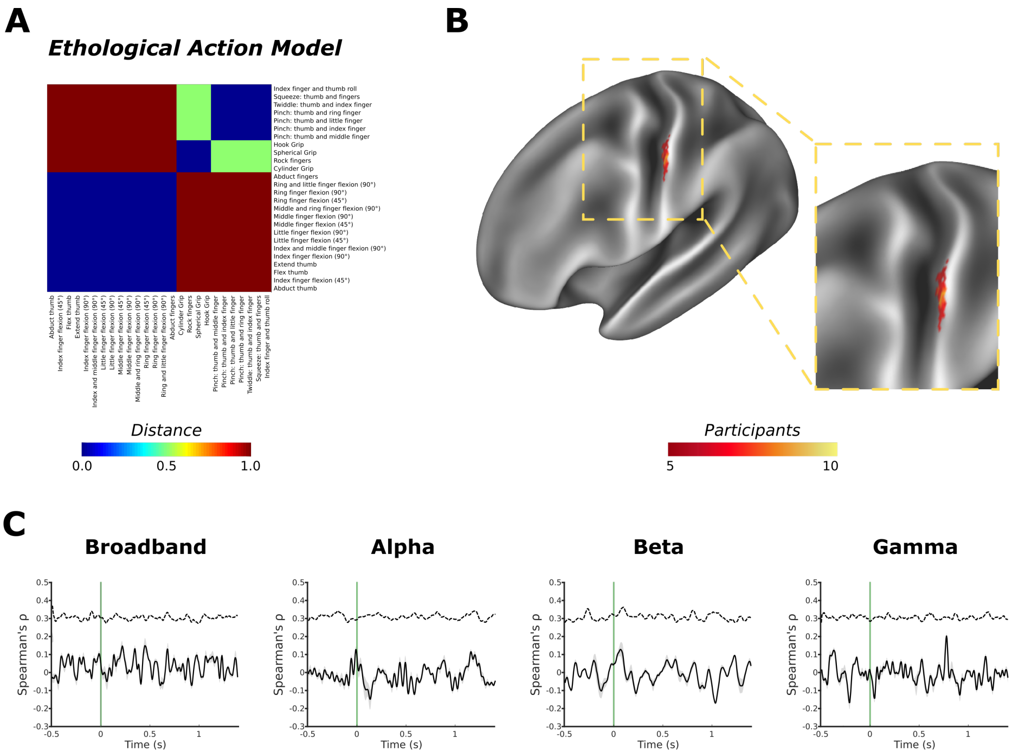


**Figure S17: Ethological action model reveals limited evidence for spatial encoding in primary somatosensory cortex.** (A) A theoretical model of cortical movement encoding on the basis of ethological action type was constructed on the basis of compelling evidence for such functional organisation in M1 from the primate literature[16]. (B) The spatial searchlight conducted using fMRI data revealed consistent cortical encoding of information in the post-central gyrus (Brodmann area 3b). (C) No evidence for the temporal encoding of this model was observed from MEG analysis.

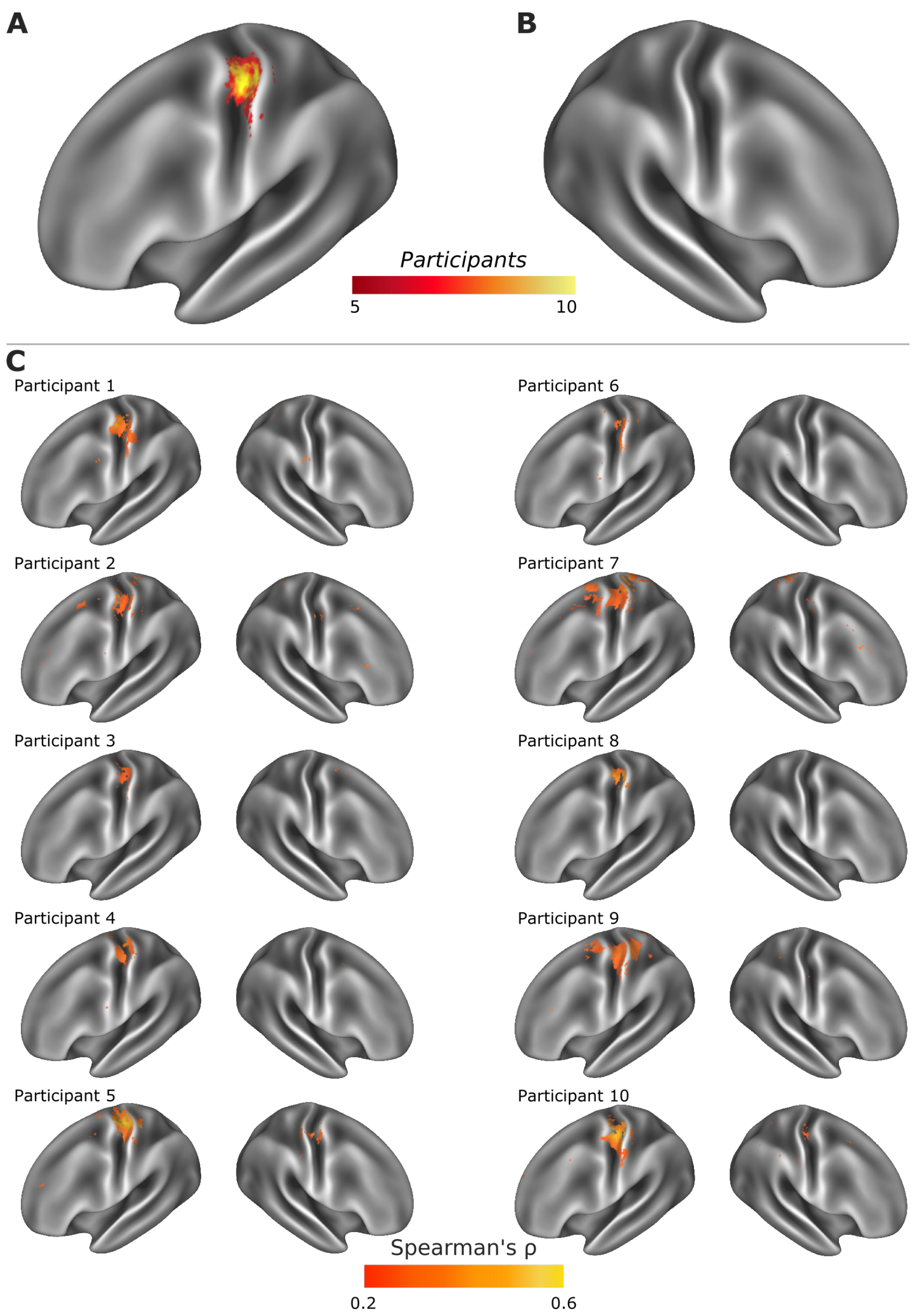


**Figure S18: fMRI searchlight analysis conducted using the kinematic model constructed from data glove recordings made during the behavioural testing session.** Evidence of the encoding of kinematic data in contralateral primary motor cortex persists using independent data glove recordings while participants were sitting upright in a more naturalistic position. Comparison with data presented in Figures 1 and 2.

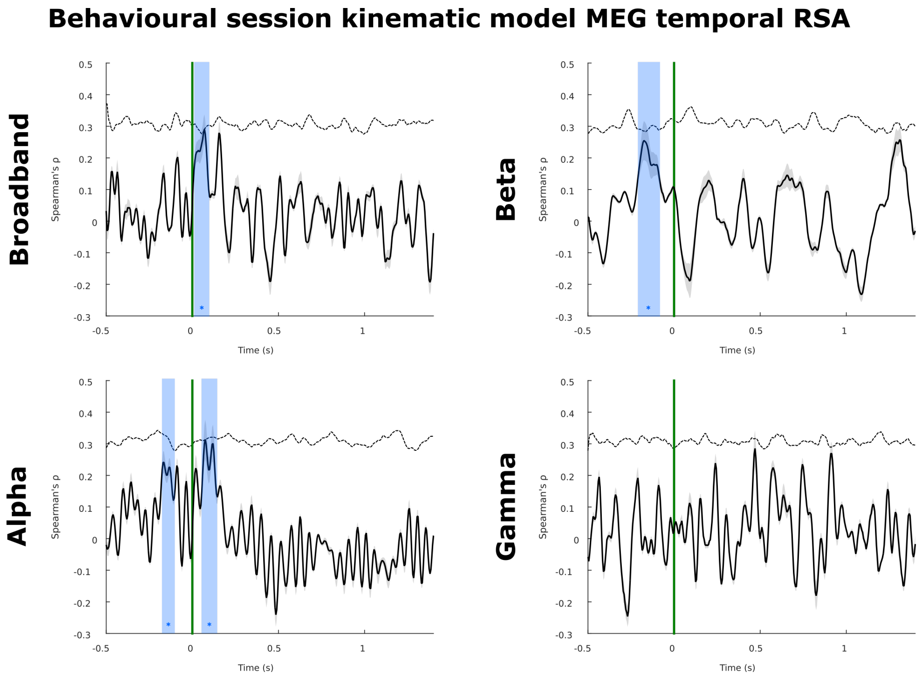


**Figure S19:** Temporal searchlight analysis investigating information encoding produces consistent results when using data glove recordings acquired during the behavioural testing session, when compared against the equivalent model derived from kinematic recordings made during the MEG sessions (Figure 3)

.
